# Environment, plant genetics, and their interaction shape important aspects of sunflower rhizosphere microbial communities

**DOI:** 10.1101/2024.08.09.607089

**Authors:** Clifton P. Bueno de Mesquita, Corinne M. Walsh, Ziv Attia, Brady D. Koehler, Zachary J. Tarble, David L. Van Tassel, Nolan C. Kane, Brent S. Hulke

## Abstract

Associations with soil microorganisms are crucial for plants’ overall health and functioning. While much work has been done to understand drivers of rhizosphere microbiome structure and function, the relative importance of geography, climate, soil properties, and plant genetics remains unclear, as results have been mixed and comprehensive studies across many sites and genotypes are limited. Rhizosphere microbiomes are crucial for crop resistance to pathogens, stress tolerance, nutrient availability, and ultimately yield. Here we quantify the relative roles of plant genotype, environment, and their interaction in shaping soil rhizosphere communities, using 16S and ITS gene sequencing of rhizosphere soils from 10 common sunflower (*Helianthus annuus*) genotypes from 15 sites across the Great Plains of the United States. While site generally outweighed genotype overall in terms of effects on archaeal, bacterial and fungal richness, community composition, and taxa relative abundances, there was also a significant interaction such that genotype exerted a significant influence on archaeal, bacterial and fungal microbiomes in certain sites. Site effects were attributed to a combination of spatial distance and differences in climate and soil properties. Microbial taxa that were previously associated with resistance to the fungal necrotrophic pathogen *Sclerotinia* were present in most sites but differed significantly in relative abundance across sites. Our results have implications for plant breeding and agronomic microbiome manipulations for agricultural improvement across different geographic regions.

**Importance:** Despite the importance of plant breeding in agriculture, we still have a limited understanding of how plant genetic variation shapes soil microbiome composition across broad geographic regions. Using 15 sites across the Great Plains of North America, we show that cultivated sunflower rhizosphere archaeal, bacterial and fungal communities are driven primarily by site soil and climatic differences, but that genotype can interact with site to influence composition, especially in warmer and drier sites with lower overall microbial richness. We also show that all taxa that were previously found to be associated with resistance to the fungal pathogen *Sclerotinia sclerotiorum* were widespread but significantly affected by site, while a subset were also significantly affected by genotype. Our results contribute to a broader understanding of rhizosphere archaeal, bacterial and fungal community assembly, and provide foundational knowledge for plant breeding efforts and potential future microbiome manipulations in agriculture.

## Introduction

Plants are no longer viewed as isolated entities, but rather as “holobionts”, which includes their aboveground and belowground associated microbiota (1, 2). Plant-associated microorganisms play a number of important roles for plant health, namely nutrient acquisition, disease resistance, and stress tolerance (3). More specifically, rhizosphere microbes can fix and cycle nitrogen, solubilize phosphorus, and produce plant-growth-promoting hormones such as auxins, gibberellins, and cytokinins (4). Therefore, microorganisms, including bacteria, archaea, fungi, protists, and other microeukaryotes, hold great potential for improving agricultural productivity. In order to successfully manipulate microbiomes to improve crop performance or breed cultivars to select for beneficial microbiomes, a comprehensive and detailed understanding of plant-associated microbial community assembly processes, including the effects of geography, climate, edaphic properties, and host plant phenotypic traits and genotype is needed.

The rhizosphere is a dynamic soil environment that is directly in contact with plant roots. While most microbes in the rhizosphere are sourced from the bulk soil (5), rhizospheres differ from bulk soils in that they are directly influenced by root exudation from the plant (3). Direct plant inputs into the rhizosphere via root exudation include a wide variety of compounds such as various sugars and carbohydrates as well as secondary metabolites and aromatic organic acids such as nicotinic, shikimic, salicylic, cinnamic and indole-3-acetic acid (6). Plant compounds can promote or inhibit microbes, and rhizosphere microbial communities have been shown to be deterministically rather than randomly selected from the bulk soil (7). However, a key outstanding question in biology is the degree to which plants actively control and select their rhizosphere and endosphere microbiomes (2). More specifically, while broader differences in rhizosphere microbial communities among plant species and functional groups have been observed (5, 8, 9), including among different plant species in the same genus (10), it is unclear to what degree different cultivars of the same plant species could select for different rhizosphere microbial communities. Such knowledge is important for agriculture, in which different crop cultivars are commonly used and new cultivars are constantly being developed, tested, and optimized. It is also relevant for the development of synthetic communities of microbes that could be applied to soils or seeds to benefit crop production (11–13).

The effects of plant genotype on rhizosphere communities have been studied in model plants such as *Arabidopsis thaliana* as well as a range of crops including maize (14, 15), rice (16, 17), soybean (18, 19), potato (20), cotton (21), sorghum (22), and sunflower (23, 24). The research has shown contrasting results. Some studies have demonstrated a strong genotype effect. For example, host genetic background and ploidy level significantly affected rhizosphere microbiota composition in *Arabidopsis thaliana* (25). However, several other studies have shown a relatively stronger effect of soil type compared to plant genotype on rhizosphere communities (18, 19, 21). A strong effect of soil type can be driven by known effects of soil properties such as pH, organic matter content, texture, cation exchange capacity, and nitrogen availability on soil microbial communities (26–28), which would then create differences in the pool of bulk soil microbes that could possibly associate with and be influenced by or be recruited by plants in the rhizosphere (5, 29). Other broader site effects can also be attributed to differences in climate (26). Purely spatial distance effects may also exist, given that rhizosphere communities have been shown to be dispersal limited (30).

The common sunflower (*Helianthus annuus*) is an important crop for a variety of products including oil, in-shell and dehulled seeds, and sprouts, and are one of the largest oilseed crops globally (31). Sunflowers provide a good opportunity to investigate the relative importance of environment versus genotype due to the wide geographic range in which they are cultivated, as well as the variety of inbred cultivars and associated genetic information that exists. Sunflower farming is currently plagued by the ascomycete fungal pathogen *Sclerotinia sclerotiorum*, which is a white rot fungus that can infect any part of the plant (32). *S. sclerotiorum* produces sclerotia which can also persist in soils (33), and plants can be initially infected through the roots. Previous work on 95 genotypes of sunflower at one site found heritable differences in rhizosphere microbiomes, and identified several bacterial operational taxonomic units (OTUs) that were associated with resistance to *S. sclerotiorum* (24), as well as several isolates from the rhizosphere that were shown to inhibit *S. sclerotiorum* growth in the laboratory (34). Because resistance to Sclerotinia basal stem rot is heritable, and correlated across environments (24), it is crucial to develop a better understanding of sunflower’s genetically-based microbial associations across multiple environments.

Here we build on this previous work by conducting a much broader survey of sunflower rhizosphere microbiomes, collecting data from 15 different sites spanning six U.S. states and 680,000 km^2^ (most of the geographic range of sunflower cultivation in North America), and 10 different genotypes selected for diversity in microbiomes and *S. sclerotiorum* resistance in previous work (24). We also assessed both archaea and bacteria (hereafter “prokaryotes”) and fungi. We tested the effects of site (and associated geographic, climatic, and edaphic differences) and plant genotype (and associated phenotypic differences) and their interaction on the rhizosphere microbiome of common sunflowers. We hypothesized that site and genotype would both significantly affect prokaryotic and fungal richness and community composition, with potentially interactive effects. We also tested the effects of site and genotype on 42 bacterial OTUs that were found to be associated with resistance to the pathogen *Sclerotinia* in a previous study of 95 genotypes at one site (24). We hypothesized that in the present multi-site dataset, there would still be significant genotype effects, but that the prevalence and relative abundances of these 42 OTUs would also vary significantly among the 15 sites.

## Methods

Rhizosphere samples were collected from 10 genotypes of sunflower at 15 different sites across six states in the U.S. Great Plains in 2020 (Table 1, Table 2). The genotypes are all highly inbred such that individuals of the same genotype can be considered genetically identical. To control for temporal effects, the sunflowers were all in the same phenophase, just before flowering, at the time of sample collection. Four samples of each genotype (two from each of two randomized, complete block replications) were collected at each site (n = 600). Samples were collected by digging up an individual plant and shaking off soil that was attached to roots into a 2 mm sieve to remove rocks and roots, and then collecting the sieved soil in a sterile 50 mL Falcon tube. An aliquot of the rhizosphere soil was then stored at −20°C for eventual DNA extraction and sequencing. Remnant soil was bulked across samples within complete block replications and subjected to soil physicochemical testing (e.g., texture, organic matter, nutrients, cations, pH; Agvise Laboratories, Northwood, ND, USA). Chlorophyll content for each sampled plant was measured just prior to rhizosphere sampling. The climate moisture index, growing degree days, mean annual temperature and precipitation and seasonality of temperature and precipitation for each site was extracted from the CHELSA database (35). DNA was extracted from leaves of each of the 10 genotypes, sequenced, and assembled as described previously to yield complete genomes for each genotype (24). Variants were called with GATK best practices (36) and only biallelic single nucleotide polymorphisms (SNPs) were retained (24). Pairwise genetic distances (Manhattan distance) among the 10 genotypes were calculated from the resulting Variant Call Format files with the *rdiversity* R package (37). Incidence of *Sclerotinia* disease in the 10 genotypes was recorded at one site (Carrington, ND, 2017), in an inoculated portion of a field with diverse crop history and multiple years of inoculated *Sclerotinia* trials. In the present study, none of the fields were inoculated with *Sclerotinia*. Additional phenotypic data were downloaded for each genotype from the U.S. Department of Agriculture’s Germplasm Resources Information Network (GRIN) database and previous work (38). Continuous variables that were available in at least 9 out of 10 genotypes were retained (days to flower, stem length, % seeds passing through 5.6 mm (14/64-inch) sieve, % seeds passing through 7.9 mm (20/64-inch) sieve, % of plants with no branching from stage R-5 to R-9, % plants resistant to *Sclerotinia* from a 2008-2009 inoculated study in North Dakota and South Dakota, weight of 100 seeds; Table 2).

**Table 1.** Site name, fertilizer and tillage management practices, sample size for 16S and ITS datasets, climate data (MAT = mean annual temperature, Tseas = temperature seasonality, MAP = mean annual precipitation, Pseas = precipitation seasonality), and selected soil data (% clay, OM = organic matter, pH, NO3- = nitrate, P = phosphorus; average of 2 replicates). The highest values of each climate and soil variable are bolded.

| Site | Management |  | n |  | Climate |  |  |  |
| --- | --- | --- | --- | --- | --- | --- | --- | --- |
|  | Fertilizer | Tillage | 16S | ITS | MAT<br>(°C) | Tseas<br>(SD*100) | MAP<br>(mm) | Pseas<br>(CV) |
| Brighton | Conventional | No-till | 40 | 39 | 9.75 | 916.25 | 391 | 50.78 |
| Brookings | Conventional | Tilled | 40 | 40 | 6.41 | 1210.95 | 591 | 61.60 |
| Burlington | Conventional | Tilled | 40 | 40 | 10.47 | 935.24 | 410 | **67.60** |
| Carrington | Conventional | Tilled | 40 | 40 | 4.49 | 1277.03 | 452 | 64.56 |
| Casselton | Conventional | Tilled | 29 | 29 | 5.17 | 1295.76 | 494 | 59.92 |
| Grandin | Conventional | Tilled | 40 | 40 | 4.84 | 1302.64 | 508 | 58.89 |
| Land Institute | Organic | Tilled | 40 | 40 | 12.95 | 1033.92 | **759** | 50.89 |
| Lindsborg-dryland | Conventional | Tilled | 40 | 40 | 12.90 | 1029.10 | 736 | 51.61 |
| Lindsborg-irrigated | Conventional | Tilled | 40 | 40 | 12.90 | 1029.10 | 736 | 51.61 |
| Mandan | Conventional | No-till | 39 | 39 | 5.75 | 1207.63 | 426 | 63.57 |
| McLaughlin | Conventional | No-till | 40 | 40 | 6.79 | 1203.40 | 425 | 64.45 |
| Mentor | Conventional | Tilled | 40 | 40 | 4.06 | **1308.66** | 575 | 63.13 |
| Pierre | Conventional | No-till | 40 | 40 | 7.62 | 1187.62 | 469 | 60.97 |
| Ralls | Conventional | Tilled | 39 | 39 | **15.23** | 833.54 | 530 | 56.23 |
| Velva | Conventional | Tilled | 40 | 40 | 4.90 | 1247.12 | 445 | 61.34 |

climate data (MAT = mean annual seasonality), and selected soil data at values of each climate and soil
| Soil |  |  |  |  |
| --- | --- | --- | --- | --- |
| Clay<br>(%) | OM<br>(%) | pH | NO<sub>3</sub><sup>-</sup><br>(ppm) | P<br>(ppm) |
| 3 | 0.3 | 5.9 | 4.75 | 10 |
| 28 | 4.7 | 7.35 | 7 | 17.5 |
| 27 | 3.3 | **7.85** | 11.5 | 18.5 |
| 16 | 3.65 | 5.7 | 3.75 | 16.5 |
| 32 | 4.35 | 6.75 | 9.25 | 22 |
| **39** | 7.3 | 6.35 | 10.5 | 38 |
| 25 | 2.65 | 7.15 | 4 | 10.5 |
| 32 | 3.85 | 5.6 | 16.75 | 17.5 |
| 15 | 2.85 | 7.7 | **33.5** | 12 |
| 25 | 4.55 | 5.4 | 8.25 | 13.5 |
| 21 | 3.25 | 5.3 | 7 | 9 |
| 21 | **10.65** | 7.2 | 15.75 | **95.5** |
| 24 | 4.1 | 6.1 | 6.75 | 10.5 |
| 11 | 0.85 | 6.25 | 8.25 | 19.5 |
| 22 | 3.6 | 5.7 | 4.5 | 14 |

**Table 2.** Genotype hierarchical clustering (Hclust, based on variant calling), pedigree identifiers, and phenotypic trait data. Phenotypic columns are chlorophyll content, days to flower, weight of 100 seeds, percent stems with no branches, stem length high range, stem length low range, percent seeds passing through a 5.6 mm (14/64 inch) sieve, percent seeds passing through a 7.9 mm (20/64 inch) sieve, incidence of Sclerotinia from the GRIN database (2008-2009 study, Taluker et al. 2014), and incidence of Sclerotinia from Carrington (2017 study).

| Hclust | Genotype | Chlorophyll | DaysToFlower | Seed wt | h | Stem |  | % Seeds pass |  |
| --- | --- | --- | --- | --- | --- | --- | --- | --- | --- |
|  |  |  |  | (g) | (%) | High | Low | 5.6 mm | 7.9 mm |
|  | PI 650839 | 19.83 | NA | 4.0 | 55 | 185 | 115 | 96 | 100 |
|  | HA 466 | 15.63 | 45 | NA | 100 | NA | NA | 36 | 100 |
|  | PI 650814 | 19.11 | 70 | 6.8 | 80 | 220 | 90 | 12 | 89 |
|  | PI 650542 | 23.47 | 56 | 6.4 | 90 | 98 | 68 | 81 | 100 |
|  | PI 650758 | 15.06 | 55 | 4.4 | 83 | 83 | 53 | 41 | 100 |
|  | PI 531361 | 20.97 | 59 | 8.0 | 50 | 180 | 130 | 26 | 100 |
|  | PI 650808 | 24.45 | 70 | 8.7 | 75 | 150 | 225 | 1 | 94 |
|  | PI 531360 | 15.34 | 63 | 7.7 | 80 | 185 | 140 | 29 | 100 |
|  | PI 650836 | 17.56 | 69 | 7.8 | 85 | 240 | 150 | 22 | 95 |
|  | PI 507903 | 17.68 | 69 | 6.3 | 40 | 180 | 85 | NA | NA |

| *Sclerotinia* |  |
| --- | --- |
| GRIN | Carr. |
| 49 | 98 |
| 24 | 21 |
| 45 | 30 |
| 25 | 12 |
| 46 | 12 |
| 16 | 7 |
| 30 | 46 |
| 34 | 43 |
| 40 | 49 |
| 27 | 58 |

DNA was extracted from 0.250 g rhizosphere soil with a Qiagen PowerSoil high throughput 96 kit (Qiagen, Hilden, Germany) following the manufacturer’s instructions. The V4 region of the 16S rRNA gene was amplified with PCR primers 515F/926R and the ITS region was amplified with PCR primers ITS1f/ITS2, following the Earth Microbiome Project protocols (39). The band size of the amplicons was checked with gel electrophoresis. PCR products were then cleaned and normalized with the SequalPrep normalization kit (Life Technologies Corporation, Carlsbad, USA). Libraries were then barcoded with 12-base Golay barcodes and pooled for sequencing. DNA sequencing was performed on an Illumina MiSeq at the BioFrontiers Institute, Boulder, CO, USA, with 2 x 250 base pair chemistry.

Raw sequence data was demultiplexed with idemp (40) and adapters were removed with cutadapt (41). Reads were then processed with the DADA2 pipeline (42) to trim and quality filter reads, infer amplicon sequence variants (ASVs), remove chimeras, and assign taxonomy with the SILVA v138.1 database for 16S (43) and UNITE v9.0 database for ITS (44). Samples were pooled for the variant calling step. Reads assigned to chloroplast, mitochondria, Eukaryota, or unassigned at the phylum level were removed from the 16S dataset with the *mctoolsr* R package (45). Reads that were not assigned to Fungi at the kingdom level or were unassigned at the phylum level were removed from the ITS dataset. Furthermore, ASVs that were extremely rare (< 15 reads across the whole dataset) were removed from both datasets. The resulting table had a mean 23784 ± 532 SE reads per sample for 16S and 23836 ± 394 SE reads per sample for ITS. ASV sequences were deposited to NCBI GenBank under BioProject ID PRJNA1114128. The filtered datasets were then normalized with center log ratio (clr) transformation and an Aitchison distance matrix was calculated with the *compositions* R package (46, 47). To complement the clr transformation and Aitchison distance analysis, ASV tables were also rarefied (48) to the lowest number for reads per sample for 16S (6027), and the second lowest number of reads per sample for ITS (4809). The total sample size for the rarefied analysis was 587 for 16S and 586 for ITS.

Bray-Curtis and Jaccard dissimilarity matrices were calculated on the rarefied 16S and ITS tables. For the 16S dataset, ASV sequences were aligned with MUSCLE (49) and a phylogenetic tree was constructed with FastTree (50). The phylogenetic tree and ASV abundance tables were used to calculate a weighted UniFrac distance matrix using the rarefied ASV table (51). Aitchison distance, weighted UniFrac distance, Jaccard dissimilarity, and Bray-Curtis dissimilarity were all strongly correlated so we just present Bray-Curtis results for interpretability. Lastly, to match previous work, bacterial and archaeal ASVs were clustered into OTUs at 97% similarity with the *DECIPHER* R package (52). Previously identified OTUs associated with genotype and *Sclerotinia* resistance (24) were identified in the current dataset with either exact sequence matching (implemented with ‘grep’) or NCBI’s Basic Local Alignment Search Tool (BLAST, identity > 99%).

The effects of site and genotype and their interaction on alpha diversity (ASV richness and Shannon diversity) were assessed with a Type III ANOVA implemented in the *car* R package (53). The effects of site and genotype and their interaction on community composition (Aitchison distance, Jaccard dissimilarity, Bray-Curtis dissimilarity, weighted UniFrac distance) were assessed with PERMANOVA implemented with the ‘adonis2’ function in the *vegan* R package (54). Site pairwise PERMANOVAs were tested with the *pairwiseAdonis* R package (55). Multivariate homogeneity of dispersions was assessed with PERMDISP implemented with the ‘betadisper’ function in *vegan*. Community composition according to Bray-Curtis dissimilarity was visualized with principal coordinates analysis (PCoA). Environmental vectors were fit to ordinations with the ‘envfit’ function in *vegan*. The effects of geographic distance, Euclidean climate distance, Euclidean soil distance, Euclidean plant phenotypic distance, and plant genetic distance on 16S and ITS community dissimilarity were assessed with both univariate and partial (controlling for geographic distance) Mantel tests implemented in *vegan*, as well as generalized dissimilarity modeling implemented in the *gdm* R package (56). Variance partitioning among spatial, climate, soil, and plant variables was performed with the ‘varpart’ function in *vegan*.

Indicator species analysis for ASV indicators of individual sites or genotypes was performed with the ‘multipatt’ function (with the ‘r.g.’ association function) in the *indicspecies* R package (57). Sets of ASVs that were shared or unique among sites and genotypes were calculated with the *MicEco* R package (58). ASV tables were summarized and plotted at different taxonomic levels using *mctoolsr*. Functional guilds were assigned with FAPROTAX for 16S (59) and FUNGuild for ITS (60). The effects of site and genotype on the relative abundance of 42 OTUs previously associated with *Sclerotinia* resistance were tested with negative binomial generalized linear models implemented in the *MASS* R package (61). All figures were made with the *ggplot2* R package (62). All analyses were performed in R version 4.2.3 (63). Data and analysis scripts are publicly available on Zenodo (doi:10.5281/zenodo.12193725).

## Results

Prokaryotic alpha diversity ranged from 569 to 2098 ASVs per sample (mean = 1500 ± 12 SE). Fungal alpha diversity ranged from 43 to 212 ASVs per sample (mean = 124 ± 1 SE). Prokaryotic ASV richness and Shannon diversity were significantly affected by site (Figure 1) but not by genotype, but there was also a significant interaction such that genotype had a significant effect in some sites but not in others (Table S1). Similarly, fungal ASV richness and Shannon diversity were significantly affected by site but not by genotype; however, there was no significant interaction (Table S1). Latitude and clay content were the top two variables that explained the most variation in prokaryotic ASV richness. Prokaryotic richness increased with increasing latitude and clay content. Climate moisture index (negative relationship) and soil phosphorus (positive), organic matter (positive), nitrate (negative), and clay content (negative) were the top predictors of fungal ASV richness (Figure S1).

**Figure 1.**
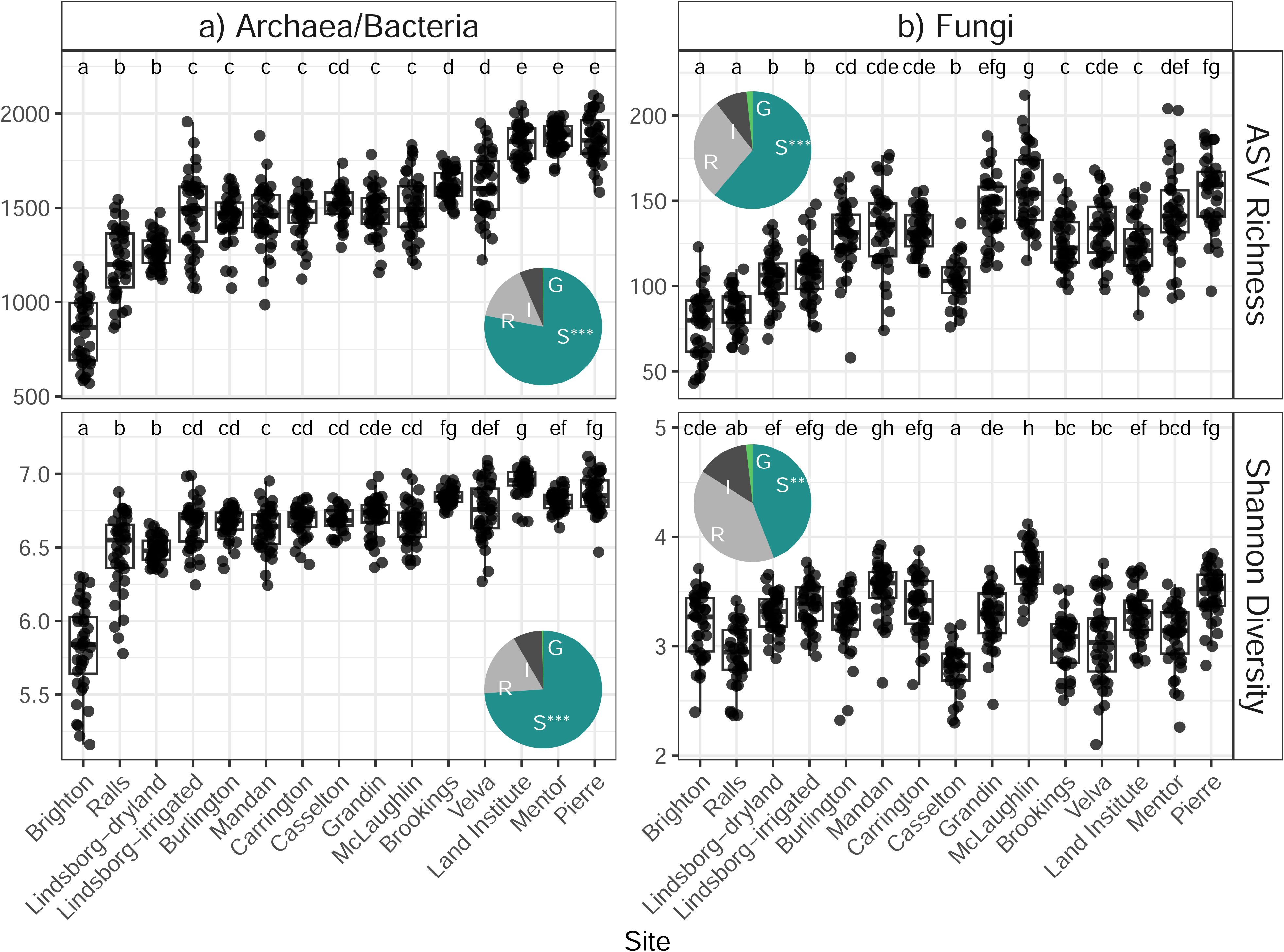
Alpha diversity (ASV richness and Shannon diversity) of prokaryotic (16S) and fungal (ITS) communities in sunflower rhizosphere soils. Insets show the effect sizes of site (S), genotype (G), the interaction term (I) and residuals (R) (*** = p < 0.001). Different letters represent significant differences among sites in each panel (Tukey posthoc p < 0.05). Note the y-axis scale varies among panels. The x-axis is sorted by overall ASV richness (prokaryotic plus fungal) and is the same in both panel columns.

Prokaryotic and fungal community composition varied significantly among sites (16S *F* = 52.9, R^2^ = 0.56, p = 0.001; ITS *F* = 69.1, R^2^ = 0.62, p = 0.001), and to a much lesser extent, among genotypes (R^2^ = 0.01, p = 0.042 and R^2^ = 0.01, p = 0.072, respectively) (Figure 2, Table S2). There was also a significant interaction between site and genotype such that genotype was more important in some sites compared to others. While restricting the PERMANOVA permutations within sites and testing the effect of genotype, genotype significantly affected both prokaryotic and fungal composition (p = 0.001), but the effect size was low (R^2^ = 0.01 for both). Most climate and soil variables were significantly associated with the principal coordinates while plant phenotype variables were not, with the exception of chlorophyll content, which was the only phenotypic variable measured at the individual level (Figure 2; envfit, p < 0.05). Pairwise PERMANOVAs demonstrated that composition at each site was significantly different than every other site (p < 0.05). Some sites had more variable prokaryotic and fungal communities than others (PERMDISP, p < 0.05), while community dispersion within genotypes was homogenous (Table S2). Sites with the most variable communities included both northern (Carrington, Grandin) and southern (Brighton, Ralls) sites. When the data were subset to individual sites, genotype significantly affected prokaryotic and fungal composition at 7 and 5 of the 15 sites, respectively. There were 3 sites in which both prokaryotes and fungi were significantly affected by genotype. On the other hand, when the data were subset to individual genotypes, sites always significantly affected microbial composition (Figure S2).

**Figure 2.**
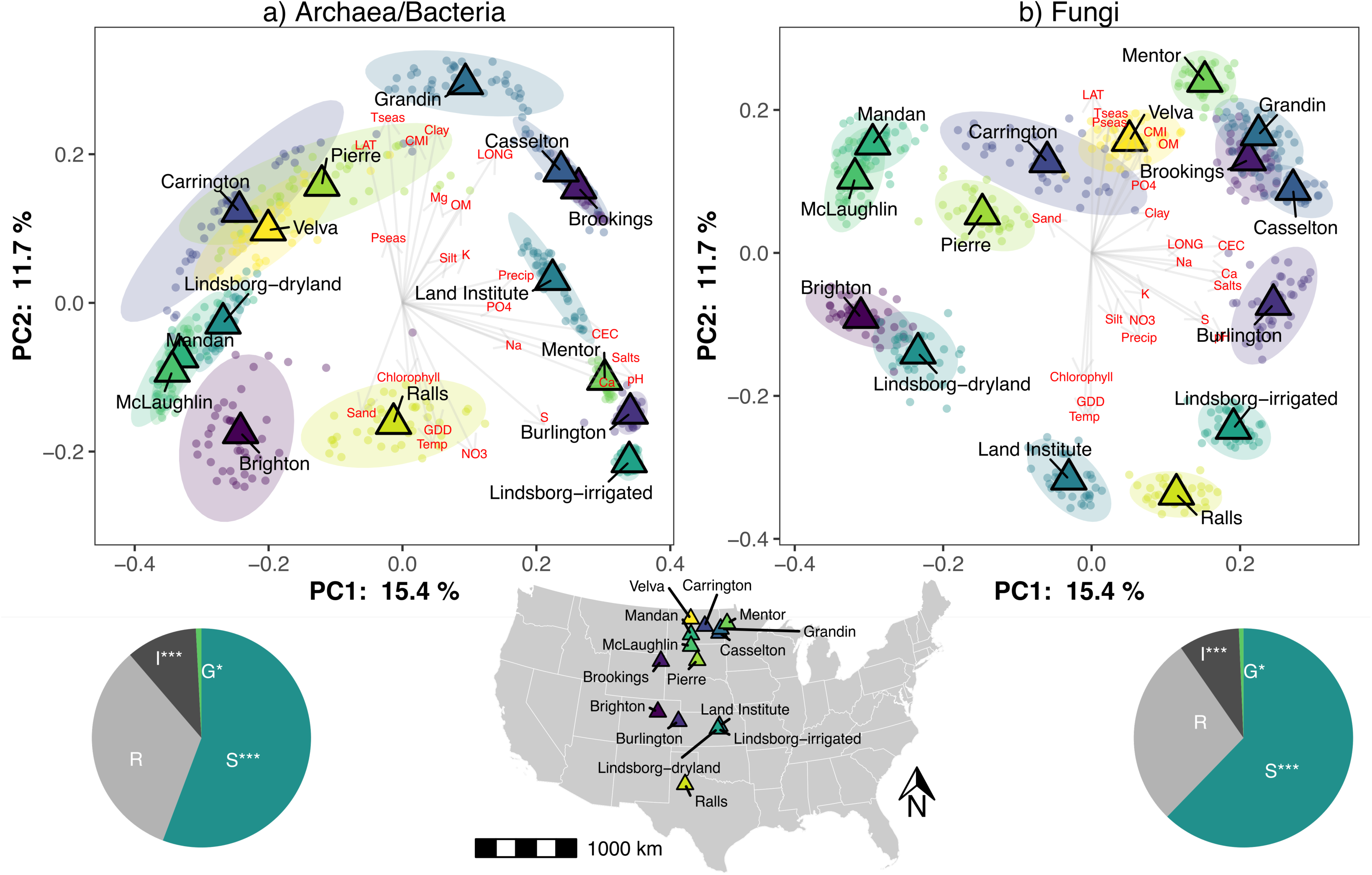
Principal coordinates analysis (PCoA) of Bray-Curtis dissimilarities for a) Prokaryotes and b) Fungi. Triangles represent site centroids. Small circles represent individual samples. Ellipses represent 95% confidence intervals around the centroid. Vectors labeled with red text represent loadings of environmental variables (GDD = growing degree days, Tseas = temperature seasonality, Pseas = precipitation seasonality, CMI = climate moisture index, LAT = latitude, LONG = longitude, OM = % organic matter, CEC = cation exchange capacity). Pies show the effect sizes of site (S), genotype (G), the interaction term (I) and residuals (R) (*** = p < 0.001). A site map is also shown.

Mantel tests, generalized dissimilarity modeling, and variance partitioning confirmed a primary effect of geography, climate, and soil properties and a secondary effect or lack of effect of plant genotype and phenotype (Table 3, Figure 3, Figure S3). Soil variables explained the most variation in prokaryote and fungal communities (20% and 23%, respectively) in the variation partitioning analysis, and had the highest maximum predicted ecological distance (MPED) values in the GDM analysis. While controlling for geographic distance, climate and soil distance were both significantly correlated with community composition, while plant phenotypic and genotypic distance were barely correlated or not correlated with community composition (Table 3). Geography and climate were tightly correlated (Mantel r = 0.96, p = 0.001) while geography and soil properties were more loosely correlated (Mantel r = 0.36, p = 0.001). The most abundant prokaryotic phyla across the whole dataset were Acidobacteriota, Proteobacteria, Planctomycetota, Actinobacteriota, Verrucomicrobiota, Chloroflexi, Bacteroidota, and Crenarchaeota, which together made up > 80% of the community (Figure 4a).

**Figure 3.**
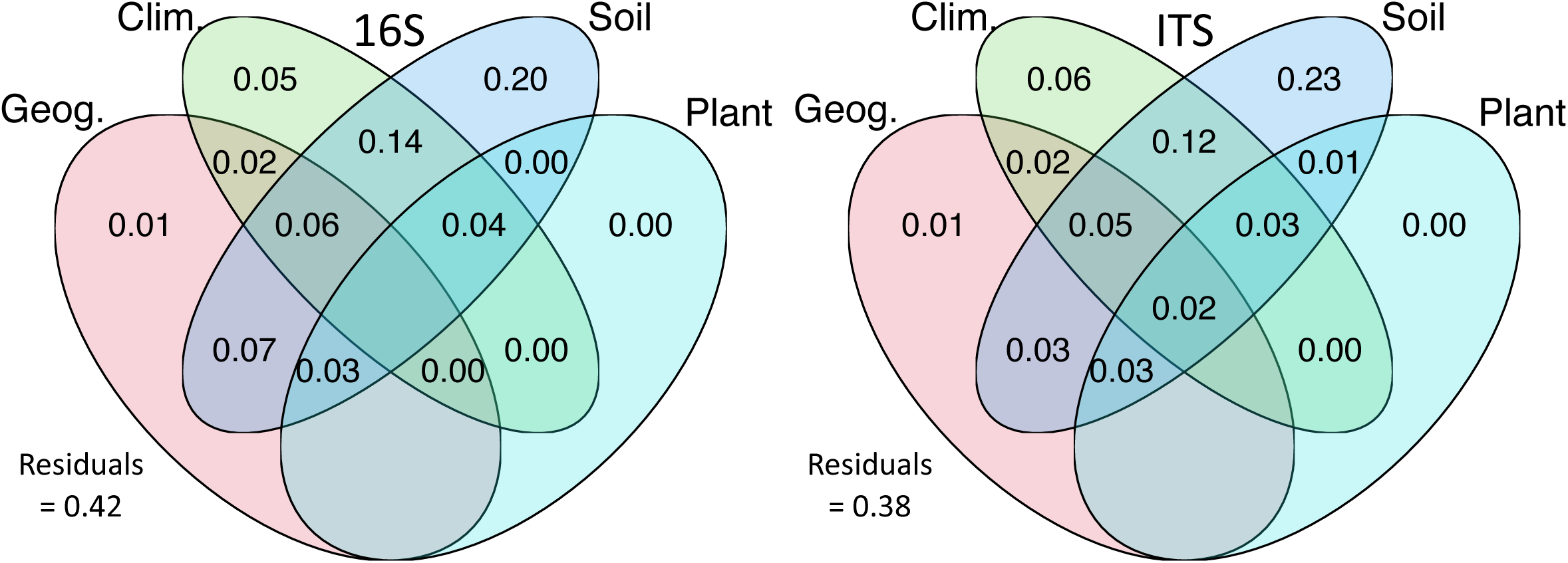
Variance partitioning of geographic, climate, soil, and plant phenotype variables for the 16S (archaea and bacteria, left) and ITS (fungi, right) datasets. Values < 0 are not shown.

**Figure 4.**
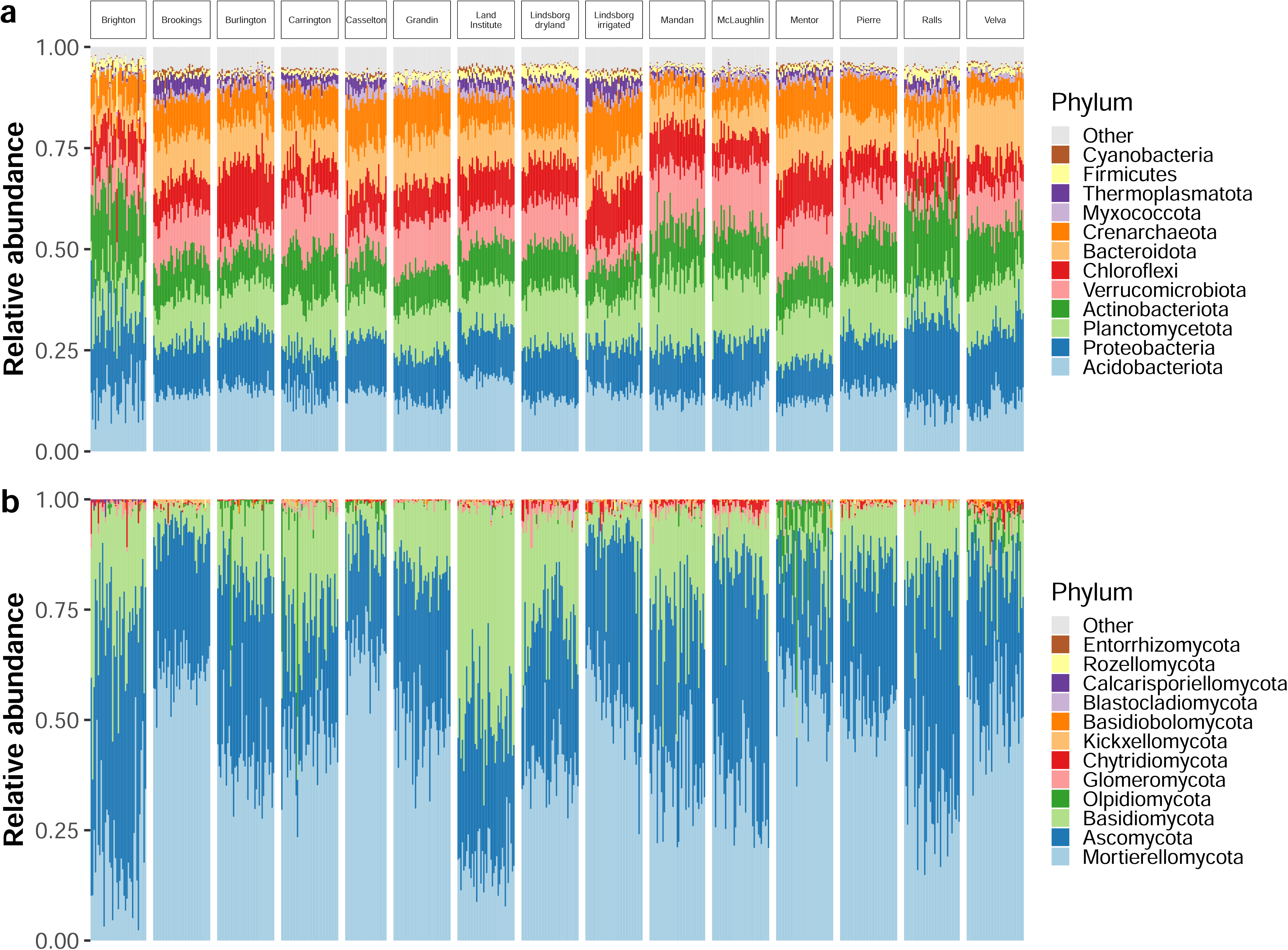
Top 12 a) archaeal and bacterial phyla and b) fungal phyla in all matching rarefied samples for each dataset (n = 586). All other phyla not in the top 12 are aggregated into the “Other category”. Phyla are sorted by overall abundance from bottom to top.

**Table 3.** Distance-based assessment of the relative effects of geographic, climate, soil, and plant variables with Mantel (geography) and partial Mantel (all other variables, controlling for geography) tests, generalized dissimilarity modeling (GDM), and variance partitioning. MPED = maximum predicted ecological distance, % = % deviance explained, VE = proportion variance explained.

| Dep. var. | Ind. var. | Mantel |  | GDM |  |  | Varpart |
| --- | --- | --- | --- | --- | --- | --- | --- |
|  |  | r | p | MPED | Retain | % | VE |
| 16S Bray-Curtis | Geography | 0.45 | 0.001 | 0.600 | Yes |  | 0.01 |
|  | Climate | 0.19 | 0.001 | 0.120 | No |  | 0.05 |
|  | Soil | **0.64** | 0.001 | **1.530** | Yes |  | **0.20** |
|  | Phenotype | -0.01 | 0.896 | 0.000 | No |  | 0.00 |
|  | Genotype | -0.01 | 0.817 | 0.010 | Yes |  | NA |
|  | Residuals |  |  |  |  | 44.03 | 0.42 |
| ITS Bray-Curtis | Geography | 0.54 | 0.001 | 0.230 | Yes |  | 0.01 |
|  | Climate | 0.33 | 0.001 | 0.561 | Yes |  | 0.06 |
|  | Soil | **0.55** | 0.001 | **0.943** | Yes |  | **0.23** |
|  | Phenotype | 0.02 | 0.041 | 0.036 | Yes |  | 0.00 |
|  | Genotype | -0.01 | 0.913 | 0.000 | No |  | NA |
|  | Residuals |  |  |  |  | 43.37 | 0.38 |

Approximately 50% of the prokaryotic reads were assigned to genus level taxonomy; the most abundant prokaryotic genera were *Candidatus Udaeobacter*, *RB41*, and *Candidatus Nitrocosmicus*, which together generally made up about 5-15% of the community (Figure S4a). The most abundant functional prokaryotic guilds were aerobic ammonia oxidizers and nitrate reducers (Figure S5a). Fungal communities were dominated (> 80%) by the Mortierellomycota, Ascomycota, and Basidiomycota phyla, although several other phyla were also detected (Figure 4b). Mortierellaceae was by far the most dominant fungal family across the whole dataset, and the two most abundant genera were from this family, *Mortierella* and *Podila*, made up ∼50% of the community in some sites (Figure S4b). Another genus of note was *Gloeoporus*, which made up ∼25% of the community at the organic Land Institute site, but was much less abundant at all other sites (Figure S4b). Most fungal taxa were classified as saprotrophs, but several other functional guilds including mycorrhizal fungi, parasites, and plant pathogens were detected.

However, a large proportion of fungal taxa were classified into multiple functional guilds (Figure S5b). In terms of prevalence, there was one bacterial ASV that was detected in all 587 samples analyzed, which was classified as belonging to the genus *Pseudarthrobacter* (Actinobacteriota). The most prevalent fungal ASVs were saprobes and plant pathogens from the Nectriaceae family, including the genera *Fusicolla*, *Fusarium*, and *Neonectria*. One ASV from the abundant *Mortierella* genus was highly prevalent (∼75% of samples). While ASVs from the *Podila* genus were highly abundant, they were not the most prevalent taxa across the whole dataset (Figure S6). There were many prokaryotic (876-2574 at p_fdr_ < 0.01) and fungal (66-259 at p_fdr_ < 0.05) ASVs associated with individual sites (Figure 5) and to a much lesser extent, each individual genotype (prokaryotes 47-147; fungi 2-15; indicator species analysis, p < 0.05, Figure S7).

**Figure 5.**
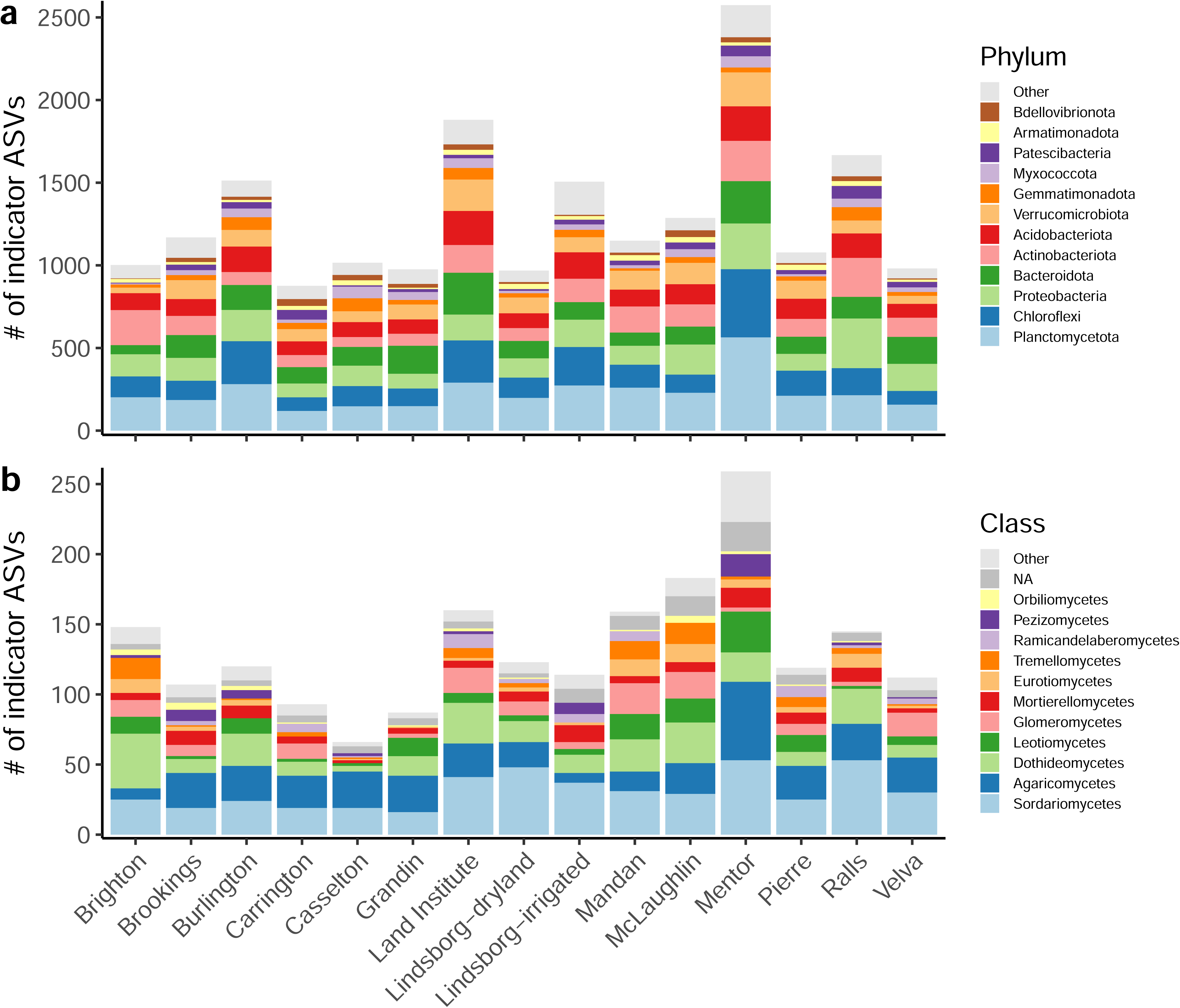
Number of site a) archaeal and bacterial indicator ASVs and b) fungal indicator ASVs, colored by prokaryotic phylum or fungal class.

Generally, site and genotype indicator ASVs were from diverse prokaryote and fungal lineages that were also some of the most abundant taxa in the dataset. Most prokaryotic site and genotype indicator ASVs were from the Planctomycetota phylum and this was consistent across all 15 sites and all 10 genotypes. Most fungal site and genotype indicator ASVs were Agaricomycetes (Basidiomycota) and Sordariomycetes (Ascomycota) (Figure S7). Mentor, Minnesota, the second most northern site in our study, was the site with consistently the most ASVs that were present only at that site and at no other sites, while Casselton, North Dakota, (120 km southwest of Mentor) consistently had the fewest site-specific ASVs. The number of ASVs found at only one site ranged from 57 (Casselton) to 1454 (Mentor) for prokaryotes and 20 (Casselton) to 262 (Mentor) for fungi. There were also 343 prokaryotic ASVs and 28 fungal ASVs present in at least one sample at all 15 sites (Figure S8). Taxonomically, these results were similar to the indicator species analysis, with Planctomycetota dominating the site-specific prokaryotic ASVs and Agaricomycetes and Sordariomycetes dominating the site-specific fungal ASVs.

All 42 OTUs identified in previous work at Carrington to be associated with *Sclerotinia* resistance were detected in the current dataset, but were grouped into 39 OTUs due to the longer reads in this dataset (generally 253 bp) compared to the original dataset (200 bp) used to calculate the 97% similarity cutoff. Three sites (Carrington, Pierre, Velva) contained all 39 of these OTUs, while the other 12 sites were missing some of these OTUs to varying degrees. At least 20 of the 39 OTUs were detected in all sites (Figure S9). Abundances of all of these OTUs varied significantly across sites, while only 11 of them were significantly affected by genotype (Figure 6, Figure S10).

**Figure 6.**
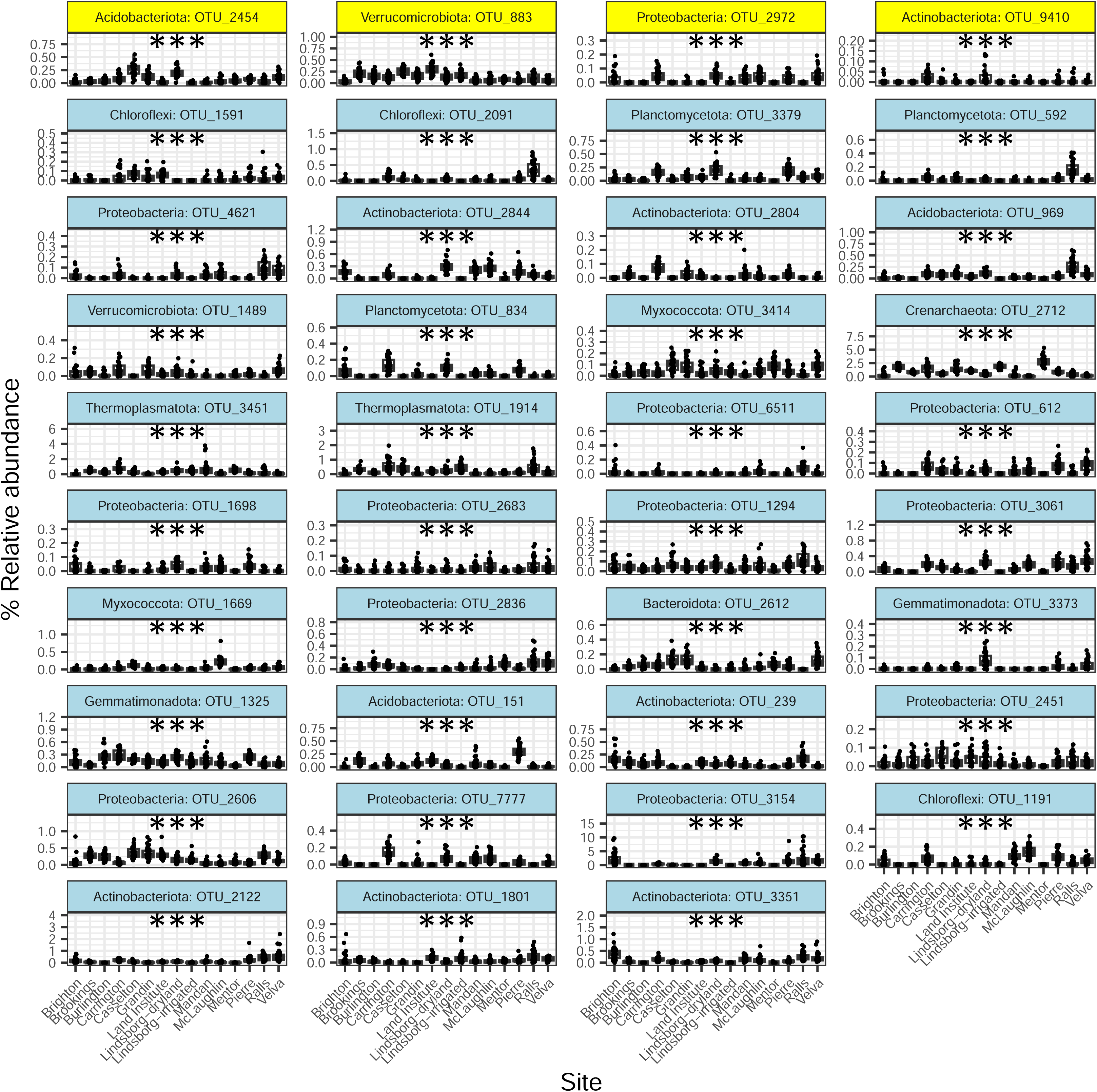
Effects of site on 39 bacterial OTUs found to be associated with *Sclerotinia* resistance in previous work at one site (Carrington). Note the y-axis scale differs among panels. Relative abundance was significantly affected by site for all 39 OTUs (*** = p < 0.001). The top row of OTUs highlighted in yellow correspond to the 4 OTUs most strongly correlated with *Sclerotinia* resistance in the previous study (24).

## Discussion

Here we used a robust sampling scheme to comprehensively assess site and sunflower genotype effects on rhizosphere microbiomes across a large geographic area of sunflower cultivation in the U.S. Great Plains, encompassing ∼1600 km in latitude and ∼650 km in longitude. Sites differed in climate as well as soil physicochemical properties (Table 1), while the 10 genotypes varied in certain phenotypic traits, including resistance to the pathogen *Sclerotinia sclerotiorum*, in addition to their known genetic differences based on whole genome sequencing and variant calling (Table 2). Four different methods – PERMANOVAs, Mantel tests, GDM, and variance partitioning – were all in agreement that site effects clearly outweighed genotype effects on both prokaryote and fungal rhizosphere communities (Figure 2, Table 3, Figure S3, Figure 3).

However, there were also significant interactions between site and genotype such that genotype increased in importance in certain sites. In particular, genotype tended to significantly affect rhizosphere communities in sites with lower ASV richness such as the warmer and drier Colorado and Texas sites (Brighton, Ralls, Figure 1, Figure S2). It is possible that in speciose sites, the speciose microbial communities overwhelm plants’ ability to more selectively control rhizosphere composition, thereby leading to no significant effects of plant genotype on composition. Alternatively, higher numbers may make it harder to detect genotypic differences with the statistical methods employed here. The degree to which different genotypes of the same plant species filter taxa from the bulk soil has been shown to vary with soil provenance (64), but the role of microbial richness has not been explicitly tested. Future work should seek to understand the role of overall soil microbial richness on plant-microbe interactions and the degree to which plant genotypes exert an influence on rhizosphere microbial communities.

Each site had a suite of ASVs that could be considered as “indicator taxa” for that site (Multipatt, p < 0.05), as well as ASVs that were found only at that site and in no others. In particular, Mentor (Minnesota) had the most unique prokaryotic and fungal ASVs. This is notable because Mentor also had the highest organic matter and highest phosphorus content, and has only recently been deforested and converted to cropland. Mentor is one of the northernmost sites in our study and the boreal forest had regrown there in the time period between prior farmland use and the current sunflower cultivation. Therefore, land use history appears to have important effects on prokaryote and fungal communities, leading to the development of unique consortia, which is consistent with the literature on land use history effects on soil microbes in general (65, 66). Individual sites also had more indicator and unique ASVs than individual genotypes, again confirming the greater effect of site on rhizosphere microbiome composition (Figure 5, Figure S7).

Differences in community composition among sites generally followed the spatial orientation of the sites, with longitude associated with PC1 and latitude associated with PC2 (Figure 2). However, there were some notable exceptions to this trend. For example, two nearby sites at Lindsborg had distinct microbial communities due to one being irrigated and one being dryland. Interestingly, the dryland site was similar to Brighton, Colorado, in prokaryotic and fungal composition, while the irrigated site was similar to Burlington, Colorado, suggesting that the irrigation management was associated with an approximately 2.4-degree shift in longitude and 19 mm of annual rainfall (Table 1). Furthermore, irrigation not only increases soil moisture levels but is also associated with changes in soil chemistry (67, 68). Indeed, in this case, the irrigated Lindsborg site had the highest soil nitrate levels of any site (Table 1).

At a broader level, plant functional group and species have been shown to have a significant influence on soil microbiomes (5, 69, 70). Finer-level effects of different plant cultivars are understudied yet could be important for agriculture, in which targeted breeding leads to the development of different cultivars of the same crop that have certain benefits. It appears that overall, especially compared to site-associated variables like climate and soil, the 10 genotypes of sunflower studied here are too similar to each other to drive major differences in rhizosphere prokaryote or fungal composition (except in certain sites, as mentioned above).

However, the relative importance of environment versus genotype could be dependent on how much environmental diversity versus genotypic diversity in the study, which is an avenue for future research and meta-analyses. Plant genetic dissimilarity was not correlated with taxonomic or phylogenetic dissimilarity in microbial communities (Table 3, Figure S2). The plant genetic dissimilarity metric in this case was made from selectively neutral genetic loci, and may not reflect dissimilarity in other specific loci that might exert more influence on microbial communities. For example, previous work at one site (Carrington) with 95 sunflower genotypes documented heritable differences in rhizosphere communities, with implications for *Sclerotinia* pathogen resistance (24). The discrepancy in genotype effects may also suggest that the ability to detect genotype effects likely depends on how much variation among the genotypes there actually is; for example, there is much more variation across 95 genotypes compared to 10 genotypes. However, in other crops, some studies have detected significant effects of only two or three genotypes (14, 18). This suggests that the effect of genotype could also depend on the crop, the degree of domestication (23), or again, how different the genotypes actually are genetically or in traits that are important for soil microbes like tissue C:N or exudate composition and production. Indeed, the degree of host plant control on rhizosphere communities has been shown to vary among crop species. For example, maize and pea have more consistent rhizosphere communities across soil types compared to onions, suggesting stronger host plant control by maize and pea plants (8).

In terms of some of the dominant prokaryotic taxa in the sunflower rhizosphere, the most abundant bacterial phylum was Acidobacteriota, despite the fact that previous work on sunflowers has shown that Acidobacteriota was more abundant in bulk soils than rhizosphere soils (71); Acidobacteriota was the third or fourth most abundant phylum in sunflower rhizospheres in South Africa (72, 73). On the other hand, the second most abundant phylum, Proteobacteria, was found to be enriched in sunflower rhizospheres compared to bulk soils (71, 72) and is typically dominant in sunflower and other plant rhizospheres (3, 72, 73). The most abundant bacterial genus across the dataset was *Candidatus Udaeobacter* (Verrucomicrobiota), which is one of the most ubiquitous and abundant soil generalists globally; it has a preference for pH ∼5, can metabolize trace gases, and is resistant to antibiotics (74–76). Its dominance in the sunflower rhizosphere could be related to the fact that *Streptomyces* (Actinobacteriota), a well-known source of antibiotics (77), was the fifth most abundant genus. The second most abundant genus, *RB41* (Acidobacteriota), has been found in high abundance in coal mining areas in China (78), and has been shown to be differentially abundant in the rhizospheres of different plant species. For example, it was more abundant in the rhizosphere of *Robinia pseudoacacia* (black locust) compared to *Pinus massoniana* (Chinese red pine) and *Cynodon dactylon* (Bermuda grass) (79). The third most abundant genus was the archaeal ammonia-oxidizer *Candidatus Nitrocosmicus* (Crenarchaeota) (80). In fact, ammonia oxidizers were one of the most abundant prokaryotic functional groups, suggesting that sunflower rhizosphere-associated taxa contribute to nitrification and the production of nitrate in these fields. There were also some other important but less abundant bacterial taxa detected in our dataset. Recent work in the laboratory using isolates have found that strains from the genera *Pseudomonas*, *Bacillus*, *Paenibacillus*, *Variovorax*, and *Sphingobacterium* can inhibit *Sclerotinia* (34) and all of these genera were present in many of our rhizosphere samples and at all 15 sites, highlighting the potential to promote native rhizosphere taxa as biocontrol agents, which is an avenue for future research.

We also built upon previous work to test the distributions of 39 OTUs that were found to be associated with *Sclerotinia* resistance (24). Such taxa are also candidates for biocontrol agents, but a better understanding of their distributions is needed first. Our work shows that the relative abundances of all 39 of these OTUs is significantly affected by site (Figure 6), while 11 of 39 are also significantly affected by genotype (Figure S10). This is again consistent with site being a more important factor than genotype. Still, the fact that 11 of these OTUs were affected by genotype means that plant breeding could still be important for promoting biocontrol agents rather than inoculation. Future work on biocontrol agents will have to take into account the environmental preferences of the taxa (81); taxa that are already ubiquitous in sunflower rhizospheres would be good potential candidates for biocontrol applications. Taxa with consistent genetically-based plant associations across environmental variation are likely to be important in affecting plant performance, disease resistance, and other traits known to be shaped by the microbiome (24), even if they are rare (82, 83). Alternatively, it is possible that in different environments the same plant genotypes form important functionally equivalent associations but with different microbial taxa. Such functional redundancy has been demonstrated in both soil and phyllosphere microbial communities (84, 85).

For fungi, *Mortierella* was the dominant genus, accounting for an astounding 25% of the community in some samples. *Mortierella* was also the most abundant genus in soybean bulk and rhizosphere soils (86). Several *Mortierella* species have been demonstrated as plant growth-promoters via different mechanisms such as increasing phosphorus or iron availability, ACC deaminase and indole-3-acetic acid production, or pathogen protection (87–90). While some studies have found increased *Mortierella* relative abundance in farms using organic fertilizer compared to inorganic fertilizer (91, 92), this was not the case in our study, as relative abundance was actually lower in our one organic site compared to the other conventional sites. The one organic site did, however, have a dramatic increase in *Gloeoporus* (Basidiomycota) relative abundance (Figure S4). *Gloeoporus* are known as wood-decay fungi and contain cosmopolitan taxa (93). In our dataset, however, *Gloeoporus* were not cosmopolitan; the genus was only present in 9% of samples, and in 8 of 15 sites. The most abundant single fungal ASV across the whole dataset was identified as *Podila*. *Podila* species appear to prefer plant-associated environments, as they have been isolated from *Pinus* and *Salix* associated soils in the subalpine, while they were not detected in higher alpine unvegetated soils (94). However, more research is needed to understand how taxa in this genus interact with plants or have any plant growth promoting potential. Saprotrophs were the most dominant fungal functional group, which is not surprising for any soil (bulk or rhizosphere), since most soil-dwelling fungi grow by consuming organic carbon. There were also some lichen and plant parasites, plant pathogens, and beneficial fungi such as arbuscular mycorrhizae and dark septate endophytes identified in the dataset, but these all had low abundances (Figure S5). The plant pathogens included *Fusarium*, which despite low abundance were some of the most ubiquitous ASVs in the dataset. *Fusarium* species are known minor pathogens of sunflowers, causing various stem wilt diseases (95–97). While the plants sampled here were all healthy, our data show that rhizosphere soils harbor *Fusarium* and that these taxa are ubiquitous across a wide geographic range.

## Conclusions

In this comprehensive survey of sunflower rhizosphere archaeal, bacterial and fungal communities across a broad geographic range spanning the Great Plains region of the U.S., we found that prokaryotic and fungal richness and community composition was mostly driven by climate and soil variables, but also are significantly shaped by sunflower genotype. Our finding that the importance of plant genotype differs across environments deserves future study. OTUs associated with *Sclerotinia* resistance (24) and genera associated with *Sclerotinia* inhibition (34) were present at most sites, but also differed significantly in relative abundance among sites. Our study shows that sunflower breeding programs could be used to manipulate microbial communities and promote beneficial microorganisms, but that site conditions will also have to be taken into account. Our study also helps highlight potential biocontrol agents and other beneficial microbes that are present across many samples and sites. For plant traits with genetically based, heritable effects that are correlated across environments, such as disease resistance, important microbial contributions to these plant traits could be due to a small subset of microbes with significant, consistent genetic effects, a hypothesis which should be confirmed by additional investigation.

## Acknowledgements

We thank Andre Gossweiler, Cameron Poyd, and Hailee Meiners for assistance with sampling rhizospheres at several sites and handling samples. We thank the Fierer Lab, especially Jessica Henley, for assistance in DNA library preparation. The authors also gratefully acknowledge the assistance of many cooperators in providing local expertise, land and field equipment, and in some cases, submitting rhizosphere samples. These include Febina Mathew, Ron Meyer, Mike Ostlie, Brian Otteson, Joel Schaefer, Tom Kirkmeyer, Karl Esping, David Archer, Jeremy Klumper, Jordan Nelson, John Swanson, Joseph Legako, and Curt Lee. We thank Cloe Pogoda, Jason Corwin, Kyle Keepers, Lara Vimercati, and Alisha Quandt for their previous analyses and work identifying potential *Sclerotinia* inhibitors. This work was supported by USDA-Agricultural Research Service CRIS projects 5442-21220-024, 3060-21220-034, 3060-21000-039, 3060-21000-043, and 3060-21000-047; and the USDA-Agricultural Research Service National *Sclerotinia* Initiative.

## References

1. Vandenkoornhuyse P, Quaiser A, Duhamel M, Le Van A, Dufresne A. 2015. The importance of the microbiome of the plant holobiont. New Phytol 206:1196–1206.

2. Sánchez-Cañizares C, Jorrín B, Poole PS, Tkacz A. 2017. Understanding the holobiont: the interdependence of plants and their microbiome. Curr Opin Microbiol 38:188–196.

3. Trivedi P, Leach JE, Tringe SG, Sa T, Singh BK. 2020. Plant–microbiome interactions: from community assembly to plant health. Nat Rev Microbiol 18:607–621.

4. Rawat P, Das S, Shankhdhar D, Shankhdhar SC. 2021. Phosphate-Solubilizing Microorganisms: Mechanism and Their Role in Phosphate Solubilization and Uptake. J Soil Sci Plant Nutr 21:49–68.

5. Berg G, Smalla K. 2009. Plant species and soil type cooperatively shape the structure and function of microbial communities in the rhizosphere. FEMS Microbiol Ecol 68:1–13.

6. Zhalnina K, Louie KB, Hao Z, Mansoori N, da Rocha UN, Shi S, Cho H, Karaoz U, Loqué D, Bowen BP, Firestone MK, Northen TR, Brodie EL. 2018. Dynamic root exudate chemistry and microbial substrate preferences drive patterns in rhizosphere microbial community assembly. Nat Microbiol 3:470–480.

7. Mendes LW, Kuramae EE, Navarrete AA, van Veen JA, Tsai SM. 2014. Taxonomical and functional microbial community selection in soybean rhizosphere. ISME J 8:1577–1587.

8. Matthews A, Pierce S, Hipperson H, Raymond B. 2019. Rhizobacterial Community Assembly Patterns Vary Between Crop Species. Front Microbiol 10.

9. Leff JW, Bardgett RD, Wilkinson A, Jackson BG, Pritchard WJ, De Long JR, Oakley S, Mason KE, Ostle NJ, Johnson D, Baggs EM, Fierer N. 2018. Predicting the structure of soil communities from plant community taxonomy, phylogeny, and traits. ISME J 12:1794– 1805.

10. Potter TS, Anacker BL, Churchill AC, Bowman WD. 2023. Plant species’ influence on rhizosphere microbial communities depends on N availability. Plant Soil 491:681–696.

11. Anušauskas J, Steponavičius D, Romaneckas K, Lekavičienė K, Zaleckas E, Sendžikienė E. 2023. The Influence of Bacteria-Inoculated Mineral Fertilizer on the Productivity and Profitability of Spring Barley Cultivation. Plants 12:1227.

12. Juknevičius D, Kriaučiūnienė Z, Jasinskas A, Šarauskis E. 2020. Analysis of Changes in Soil Organic Carbon, Energy Consumption and Environmental Impact Using Bio-Products in the Production of Winter Wheat and Oilseed Rape. 19. Sustainability 12:8246.

13. Baas P, Bell C, Mancini LM, Lee MN, Conant RT, Wallenstein MD. 2016. Phosphorus mobilizing consortium Mammoth P^TM^ enhances plant growth. PeerJ 4:e2121.

14. Aira M, Gómez-Brandón M, Lazcano C, Bååth E, Domínguez J. 2010. Plant genotype strongly modifies the structure and growth of maize rhizosphere microbial communities. Soil Biol Biochem 42:2276–2281.

15. Peiffer JA, Spor A, Koren O, Jin Z, Tringe SG, Dangl JL, Buckler ES, Ley RE. 2013. Diversity and heritability of the maize rhizosphere microbiome under field conditions. Proc Natl Acad Sci 110:6548–6553.

16. Xiong J, Lu J, Li X, Qiu Q, Chen J, Yan C. 2021. Effect of rice (Oryza sativa L.) genotype on yield: Evidence from recruiting spatially consistent rhizosphere microbiome. Soil Biol Biochem 161:108395.

17. Shenton M, Iwamoto C, Kurata N, Ikeo K. 2016. Effect of Wild and Cultivated Rice Genotypes on Rhizosphere Bacterial Community Composition. Rice 9:42.

18. Xu Y, Wang G, Jin J, Liu J, Zhang Q, Liu X. 2009. Bacterial communities in soybean rhizosphere in response to soil type, soybean genotype, and their growth stage. Soil Biol Biochem 41:919–925.

19. Liu F, Hewezi T, Lebeis SL, Pantalone V, Grewal PS, Staton ME. 2019. Soil indigenous microbiome and plant genotypes cooperatively modify soybean rhizosphere microbiome assembly. BMC Microbiol 19:201.

20. Brolsma KM, Vonk JA, Mommer L, Van Ruijven J, Hoffland E, De Goede RGM. 2017. Microbial catabolic diversity in and beyond the rhizosphere of plant species and plant genotypes. Pedobiologia 61:43–49.

21. Yang C, Yue H, Ma Z, Feng Z, Feng H, Zhao L, Zhang Y, Deakin G, Xu X, Zhu H, Wei F. 2022. Influence of plant genotype and soil on the cotton rhizosphere microbiome. Front Microbiol 13.

22. Deng S, Caddell DF, Xu G, Dahlen L, Washington L, Yang J, Coleman-Derr D. 2021. Genome wide association study reveals plant loci controlling heritability of the rhizosphere microbiome. ISME J 15:3181–3194.

23. Leff JW, Lynch RC, Kane NC, Fierer N. 2017. Plant domestication and the assembly of bacterial and fungal communities associated with strains of the common sunflower, Helianthus annuus. New Phytol 214:412–423.

24. Pogoda CS, Keepers KG, Reinert S, Talukder ZI, Smart BC, Attia Z, Corwin JA, Money KL, Collier-zans ECE, Underwood W, Gulya TJ, Quandt CA, Kane NC, Hulke BS. 2024. Heritable differences in abundance of bacterial rhizosphere taxa are correlated with fungal necrotrophic pathogen resistance. Mol Ecol 33:e17218.

25. Ponsford JCB, Hubbard CJ, Harrison JG, Maignien L, Buerkle CA, Weinig C. 2022. Whole-Genome Duplication and Host Genotype Affect Rhizosphere Microbial Communities. mSystems 7:e00973–21.

26. Nuccio EE, Anderson-Furgeson J, Estera KY, Pett-Ridge J, de Valpine P, Brodie EL, Firestone MK. 2016. Climate and edaphic controllers influence rhizosphere community assembly for a wild annual grass. Ecology 97:1307–1318.

27. Lauber CL, Hamady M, Knight R, Fierer N. 2009. Pyrosequencing-Based Assessment of Soil pH as a Predictor of Soil Bacterial Community Structure at the Continental Scale. Appl Environ Microbiol 75:5111–5120.

28. King AJ, Freeman KR, McCormick KF, Lynch RC, Lozupone C, Knight R, Schmidt SK. 2010. Biogeography and habitat modelling of high-alpine bacteria. Nat Commun 1:53.

29. Bell CW, Asao S, Calderon F, Wolk B, Wallenstein MD. 2015. Plant nitrogen uptake drives rhizosphere bacterial community assembly during plant growth. Soil Biol Biochem 85:170– 182.

30. Zhang G, Wei G, Wei F, Chen Z, He M, Jiao S, Wang Y, Dong L, Chen S. 2021. Dispersal Limitation Plays Stronger Role in the Community Assembly of Fungi Relative to Bacteria in Rhizosphere Across the Arable Area of Medicinal Plant. Front Microbiol 12.

31. Adeleke BS, Babalola OO. 2020. Oilseed crop sunflower (Helianthus annuus) as a source of food: Nutritional and health benefits. Food Sci Nutr 8:4666–4684.

32. Hegedus DD, Rimmer SR. 2005. Sclerotinia sclerotiorum: When “to be or not to be” a pathogen? FEMS Microbiol Lett 251:177–184.

33. Erental A, Dickman MB, Yarden O. 2008. Sclerotial development in Sclerotinia sclerotiorum: awakening molecular analysis of a “Dormant” structure. Fungal Biol Rev 22:6–16.

34. Vimercati L, Bueno de Mesquita C, Kane N, Hulke B, Quandt C. 2024. Bacterial inhibitors of the fungal phytopathogen Sclerotinia sclerotiorum isolated from rhizosphere soils of resistant sunflower genotypes. Plant Dis In preparation.

35. Karger DN, Conrad O, Böhner J, Kawohl T, Kreft H, Soria-Auza RW, Zimmermann NE, Linder HP, Kessler M. 2017. Climatologies at high resolution for the earth’s land surface areas. Sci Data 4:170122.

36. der Auwera GAV, O’Connor BD. 2020. Genomics in the Cloud: Using Docker, GATK, and WDL in Terra. O’Reilly Media, Inc.

37. Mitchell S, Reeve R, White T. 2022. rdiversity. R package version 2.2. https://github.com/boydorr/rdiversity (2.2). R.

38. Talukder ZI, Hulke BS, Marek LF, Gulya TJ. 2014. Sources of Resistance to Sunflower Diseases in a Global Collection of Domesticated USDA Plant Introductions. Crop Sci 54:694–705.

39. Thompson LR, Sanders JG, McDonald D, Amir A, Ladau J, Locey KJ, Prill RJ, Tripathi A, Gibbons SM, Ackermann G, Navas-Molina JA, Janssen S, Kopylova E, Vázquez-Baeza Y, González A, Morton JT, Mirarab S, Zech Xu Z, Jiang L, Haroon MF, Kanbar J, Zhu Q, Jin Song S, Kosciolek T, Bokulich NA, Lefler J, Brislawn CJ, Humphrey G, Owens SM, Hampton-Marcell J, Berg-Lyons D, McKenzie V, Fierer N, Fuhrman JA, Clauset A, Stevens RL, Shade A, Pollard KS, Goodwin KD, Jansson JK, Gilbert JA, Knight R. 2017. A communal catalogue reveals Earth’s multiscale microbial diversity. Nature 551:457–463.

40. Wu Y. 2014. idemp.

41. Martin M. 2011. Cutadapt removes adapter sequences from high-throughput sequencing reads. 1. EMBnet.journal 17:10–12.

42. Callahan BJ, McMurdie PJ, Rosen MJ, Han AW, Johnson AJA, Holmes SP. 2016. DADA2: High-resolution sample inference from Illumina amplicon data. Nat Methods 13:581–583.

43. Quast C, Pruesse E, Yilmaz P, Gerken J, Schweer T, Yarza P, Peplies J, Glöckner FO. 2013. The SILVA ribosomal RNA gene database project: improved data processing and web-based tools. Nucleic Acids Res 41:D590–D596.

44. Abarenkov K, Nilsson RH, Larsson K-H, Taylor AFS, May TW, Frøslev TG, Pawlowska J, Lindahl B, Põldmaa K, Truong C, Vu D, Hosoya T, Niskanen T, Piirmann T, Ivanov F, Zirk A, Peterson M, Cheeke TE, Ishigami Y, Jansson AT, Jeppesen TS, Kristiansson E, Mikryukov V, Miller JT, Oono R, Ossandon FJ, Paupério J, Saar I, Schigel D, Suija A, Tedersoo L, Kõljalg U. 2024. The UNITE database for molecular identification and taxonomic communication of fungi and other eukaryotes: sequences, taxa and classifications reconsidered. Nucleic Acids Res 52:D791–D797.

45. Leff JW. 2022. mctoolsr: Microbial Community Data Analysis Tools (0.1.1.9). R.

46. van den Boogaart K, Tolosana-Delgado R, Bren M. 2023. compositions: Compositional Data Analysis (2.0-5).

47. Gloor GB, Macklaim JM, Pawlowsky-Glahn V, Egozcue JJ. 2017. Microbiome Datasets Are Compositional: And This Is Not Optional. Front Microbiol 8.

48. Schloss PD. 2024. Rarefaction is currently the best approach to control for uneven sequencing effort in amplicon sequence analyses. mSphere 0:e00354–23.

49. Edgar RC. 2004. MUSCLE: multiple sequence alignment with high accuracy and high throughput. Nucleic Acids Res 32:1792–1797.

50. Price MN, Dehal PS, Arkin AP. 2009. FastTree: Computing Large Minimum Evolution Trees with Profiles instead of a Distance Matrix. Mol Biol Evol 26:1641–1650.

51. Lozupone CA, Hamady M, Kelley ST, Knight R. 2007. Quantitative and Qualitative β Diversity Measures Lead to Different Insights into Factors That Structure Microbial Communities. Appl Environ Microbiol 73:1576–1585.

52. Wright E S. 2016. Using DECIPHER v2.0 to Analyze Big Biological Sequence Data in R. R J 8:352.

53. Fox J, Weisberg S. 2018. An R Companion to Applied Regression, 3rd ed. SAGE Publications, Thousand Oaks, CA.

54. Oksanen J, Blanchet FG, Friendly M, Kindt R, Legendre P, McGlinn D, Minchin PR, O’Hara RB, Simpson GL, Solymos P, Stevens MHH, Szoecs E, Wagner H. 2022. vegan: Community Ecology Package. R package version 2.5–6. https://CRAN.R-project.org/package=vegan (2.6-4). R.

55. Martinez Arbizu P. 2017. pairwiseAdonis: Pairwise Multilevel Comparison using Adonis (R package version 0.4.1).

56. Fitzpatrick M, Mokany K, Manion G, Nieto-Lugilde D, Ferrier S. 2022. gdm: Generalized Dissimilarity Modeling (R package version 1.5.0-9.1). R.

57. De Cáceres M, Legendre P. 2009. Associations between species and groups of sites: indices and statistical inference. Ecology 90:3566–3574.

58. Russel J. 2024. MicEco: Various functions for microbial community data. R package version 0.9.19. (0.9.19). R.

59. Louca S, Parfrey LW, Doebeli M. 2016. Decoupling function and taxonomy in the global ocean microbiome. Science 353:1272–1277.

60. Nguyen NH, Song Z, Bates ST, Branco S, Tedersoo L, Menke J, Schilling JS, Kennedy PG. 2016. FUNGuild: An open annotation tool for parsing fungal community datasets by ecological guild. Fungal Ecol 20:241–248.

61. Venables W, Ripley B. 2002. Modern Applied Statistics with SFourth Edition. Springer, New York, NY.

62. Wickham H. 2016. ggplot2: Elegant Graphics for Data Analysis. Springer-Verlag, New York, NY. https://ggplot2.tidyverse.org.

63. R Core Team. 2023. R: A Language and Environment for Statistical Computing (4.2.3). R. R Foundation for Statistical Computing, Vienna, Austria.

64. Meade CV, Bueno de Mesquita CP, Schmidt SK, Suding KN. 2020. The presence of a foreign microbial community promotes plant growth and reduces filtering of root fungi in the arctic-alpine plant Silene acaulis. Plant Ecol Divers 13:377–390.

65. Jangid K, Williams MA, Franzluebbers AJ, Schmidt TM, Coleman DC, Whitman WB. 2011. Land-use history has a stronger impact on soil microbial community composition than aboveground vegetation and soil properties. Soil Biol Biochem 43:2184–2193.

66. Osburn ED, Aylward FO, Barrett JE. 2021. Historical land use has long-term effects on microbial community assembly processes in forest soils. ISME Commun 1:48.

67. Amundson RG, Smith VS. 1988. Effects of irrigation on the chemical properties of a soil in the western san joaquin valley, California. Arid Soil Res Rehabil 2:1–17.

68. Modupe A O. O, Olabiyi O. 2014. Effects of Irrigation Practices on Some Soil Chemical Properties on OMI Irrigation Scheme. Int J Eng Res Appl 4:29–35.

69. Garbeva P, van Elsas JD, van Veen JA. 2008. Rhizosphere microbial community and its response to plant species and soil history. Plant Soil 302:19–32.

70. Dassen S, Cortois R, Martens H, de Hollander M, Kowalchuk GA, van der Putten WH, De Deyn GB. 2017. Differential responses of soil bacteria, fungi, archaea and protists to plant species richness and plant functional group identity. Mol Ecol 26:4085–4098.

71. Nwachukwu BC, Ayangbenro AS, Babalola OO. 2023. Structural diversity of bacterial communities in two divergent sunflower rhizosphere soils. Ann Microbiol 73:9.

72. Oberholster T, Vikram S, Cowan D, Valverde A. 2018. Key microbial taxa in the rhizosphere of sorghum and sunflower grown in crop rotation. Sci Total Environ 624:530–539.

73. Alawiye TT, Babalola OO. 2021. Metagenomic Insight into the Community Structure and Functional Genes in the Sunflower Rhizosphere Microbiome. 2. Agriculture 11:167.

74. Brewer TE, Handley KM, Carini P, Gilbert JA, Fierer N. 2016. Genome reduction in an abundant and ubiquitous soil bacterium ‘Candidatus Udaeobacter copiosus.’ Nat Microbiol 2:1–7.

75. Hofer U. 2016. A small soil bacterium dominates. Nat Rev Microbiol 14:729–729.

76. Willms IM, Bolz SH, Yuan J, Krafft L, Schneider D, Schöning I, Schrumpf M, Nacke H. 2021. The ubiquitous soil verrucomicrobial clade ‘Candidatus Udaeobacter’ shows preferences for acidic pH. Environ Microbiol Rep 13:878–883.

77. Chater KF. 2016. Recent advances in understanding Streptomyces. F1000Research 5:2795.

78. Sun J, Yang L, Wei J, Quan J, Yang X. 2020. The responses of soil bacterial communities and enzyme activities to the edaphic properties of coal mining areas in Central China. PLOS ONE 15:e0231198.

79. Liu C, Zhuang J, Wang J, Fan G, Feng M, Zhang S. 2022. Soil bacterial communities of three types of plants from ecological restoration areas and plant-growth promotional benefits of Microbacterium invictum (strain X-18). Front Microbiol 13.

80. Lehtovirta-Morley LE, Ross J, Hink L, Weber EB, Gubry-Rangin C, Thion C, Prosser JI, Nicol GW. 2016. Isolation of ‘Candidatus Nitrosocosmicus franklandus’, a novel ureolytic soil archaeal ammonia oxidiser with tolerance to high ammonia concentration. FEMS Microbiol Ecol 92:fiw057.

81. Leggett M, Leland J, Kellar K, Epp B. 2011. Formulation of microbial biocontrol agents – an industrial perspective. Can J Plant Pathol 33:101–107.

82. Chen Q-L, Ding J, Zhu D, Hu H-W, Delgado-Baquerizo M, Ma Y-B, He J-Z, Zhu Y-G. 2020. Rare microbial taxa as the major drivers of ecosystem multifunctionality in long-term fertilized soils. Soil Biol Biochem 141:107686.

83. Zhang Y, Dong S, Gao Q, Ganjurjav H, Wang X, Geng W. 2019. “Rare biosphere” plays important roles in regulating soil available nitrogen and plant biomass in alpine grassland ecosystems under climate changes. Agric Ecosyst Environ 279:187–193.

84. Chen H, Ma K, Lu C, Fu Q, Qiu Y, Zhao J, Huang Y, Yang Y, Schadt CW, Chen H. 2022. Functional Redundancy in Soil Microbial Community Based on Metagenomics Across the Globe. Front Microbiol 13.

85. Lambais MR, Barrera SE, Santos EC, Crowley DE, Jumpponen A. 2017. Phyllosphere Metaproteomes of Trees from the Brazilian Atlantic Forest Show High Levels of Functional Redundancy. Microb Ecol 73:123–134.

86. Liu J, Yao Q, Li Y, Zhang W, Mi G, Chen X, Yu Z, Wang G. 2019. Continuous cropping of soybean alters the bulk and rhizospheric soil fungal communities in a Mollisol of Northeast PR China. Land Degrad Dev 30:1725–1738.

87. Ozimek E, Hanaka A. 2021. Mortierella Species as the Plant Growth-Promoting Fungi Present in the Agricultural Soils. 1. Agriculture 11:7.

88. Li F, Zhang S, Wang Y, Li Y, Li P, Chen L, Jie X, Hu D, Feng B, Yue K, Han Y. 2020. Rare fungus, Mortierella capitata, promotes crop growth by stimulating primary metabolisms related genes and reshaping rhizosphere bacterial community. Soil Biol Biochem 151:108017.

89. Wang Y, Wang L, Suo M, Qiu Z, Wu H, Zhao M, Yang H. 2022. Regulating Root Fungal Community Using Mortierella alpina for Fusarium oxysporum Resistance in Panax ginseng. Front Microbiol 13.

90. Sang Y, Jin L, Zhu R, Yu X-Y, Hu S, Wang B-T, Ruan H-H, Jin F-J, Lee H-G. 2022. Phosphorus-Solubilizing Capacity of Mortierella Species Isolated from Rhizosphere Soil of a Poplar Plantation. 12. Microorganisms 10:2361.

91. Qiu W, Su H, Yan L, Ji K, Liu Q, Jian H. 2020. Organic Fertilization Assembles Fungal Communities of Wheat Rhizosphere Soil and Suppresses the Population Growth of Heterodera avenae in the Field. Front Plant Sci 11.

92. Li F, Chen L, Redmile-Gordon M, Zhang J, Zhang C, Ning Q, Li W. 2018. Mortierella elongata’s roles in organic agriculture and crop growth promotion in a mineral soil. Land Degrad Dev 29:1642–1651.

93. Cho Y, Seo CW, Jung PE, Lim YW. 2023. Global phylogeographical distribution of Gloeoporus dichrous. PLOS ONE 18:e0288498.

94. Telagathoti A, Probst M, Peintner U. 2021. Habitat, Snow-Cover and Soil pH, Affect the Distribution and Diversity of Mortierellaceae Species and Their Associations to Bacteria. Front Microbiol 12.

95. Haggag WM,. AWA. 2001. Efficiency of Trichoderma Species on Control of Fusarium-rot, Root Knot and Reniform Nematodes Disease Complex on Sunflower. Pak J Biol Sci 4:314– 318.

96. Sattarovich SB, Normamadovich RU, Kakhramonovich KU, Mirodilovich AM. 2020. FUNGAL DISEASES OF SUNFLOWER AND MEASURES AGAINST THEM. 6. PalArchs J Archaeol Egypt Egyptol 17:3268–3279.

97. Gontcharov SV, Antonova TS, Saukova SL. 2006. Sunflower Breeding For Resistance To Fusarium / Selección De Girasol Por Resistencia A Fusarium / Sélection De Tournesol Pour La Résistance Au Fongus Fusarium. Helia 29:49–54.

