## Supplemental Figures for "Environment, plant genetics, and their interaction shape important aspects of sunflower rhizosphere microbial communities"

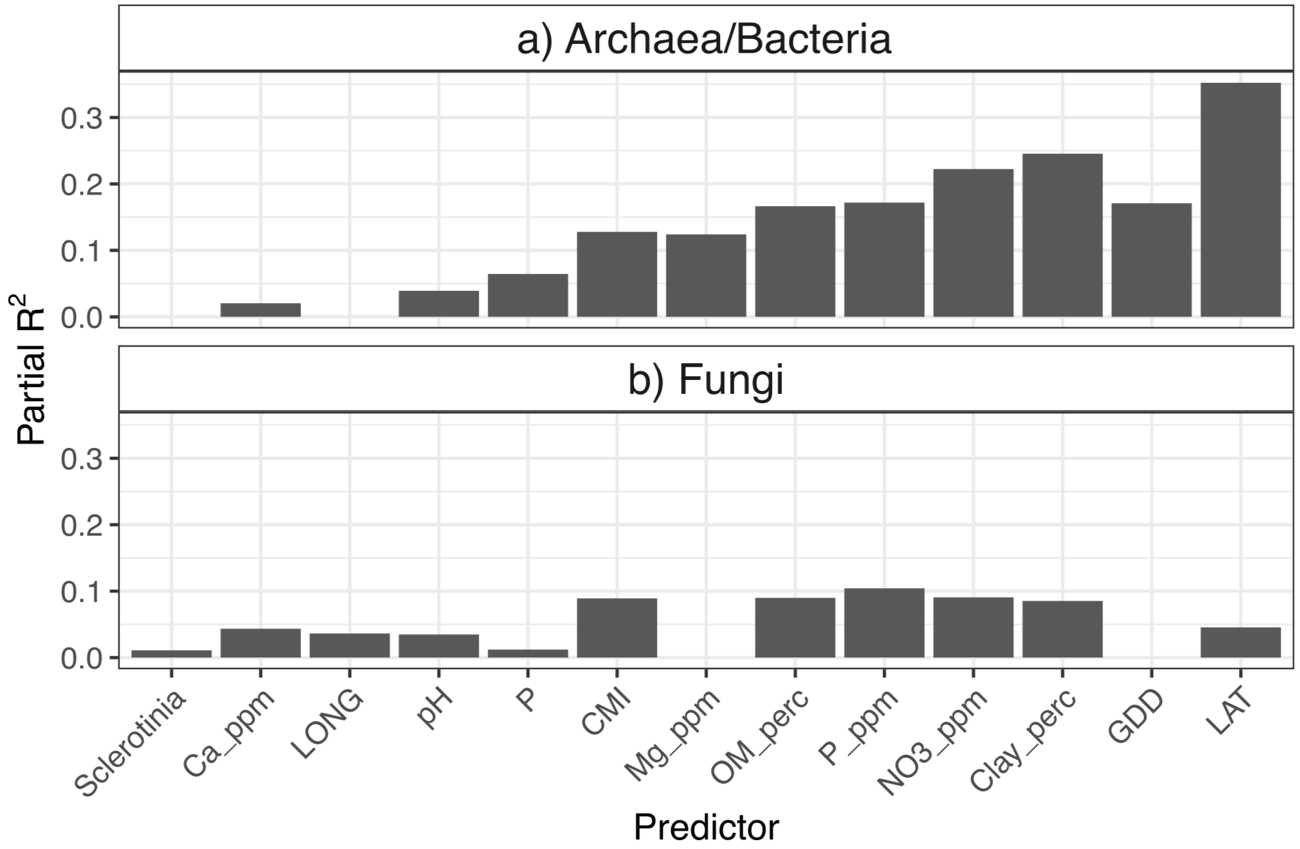

Figure S1. Partial R^2^ values for best models predicting a) archaeal/bacteria and b) fungal ASV richness in the rarefied datasets. The x-axis is sorted by increasing combined partial R^2^. Note that *Sclerotinia* resistance and longitude were not included in the best model of prokaryotic ASV richness, while Mg and GDD were not included in the best model of fungal ASV richness.

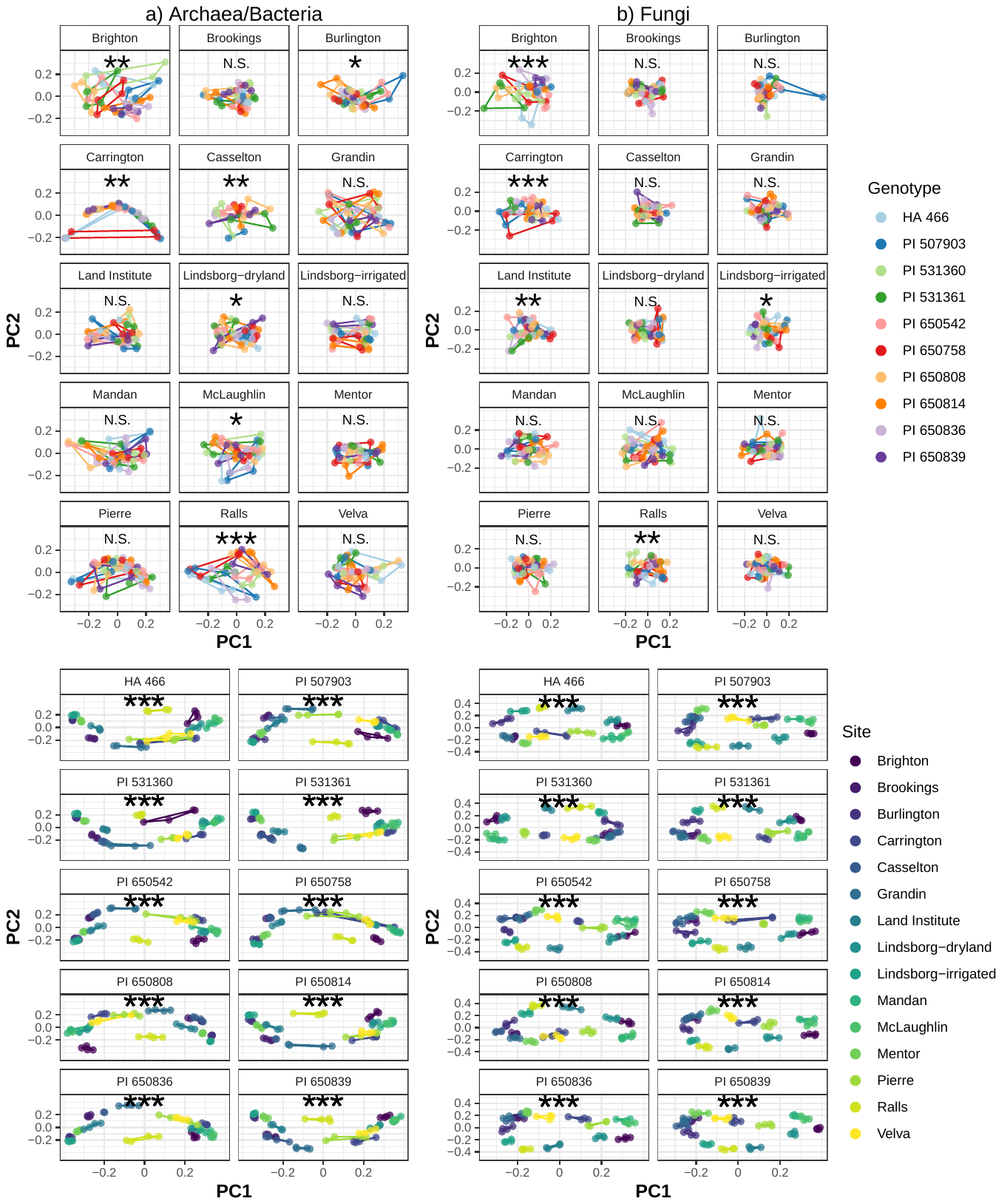

Figure S2. Effects of site when the dataset was subset to each genotype and effects of genotype when the dataset was subset to each site, for a) Archaea/Bacteria and b) Fungi. *** = p < 0.001, ** p < 0.01, * p < 0.05 from PERMANOVA.

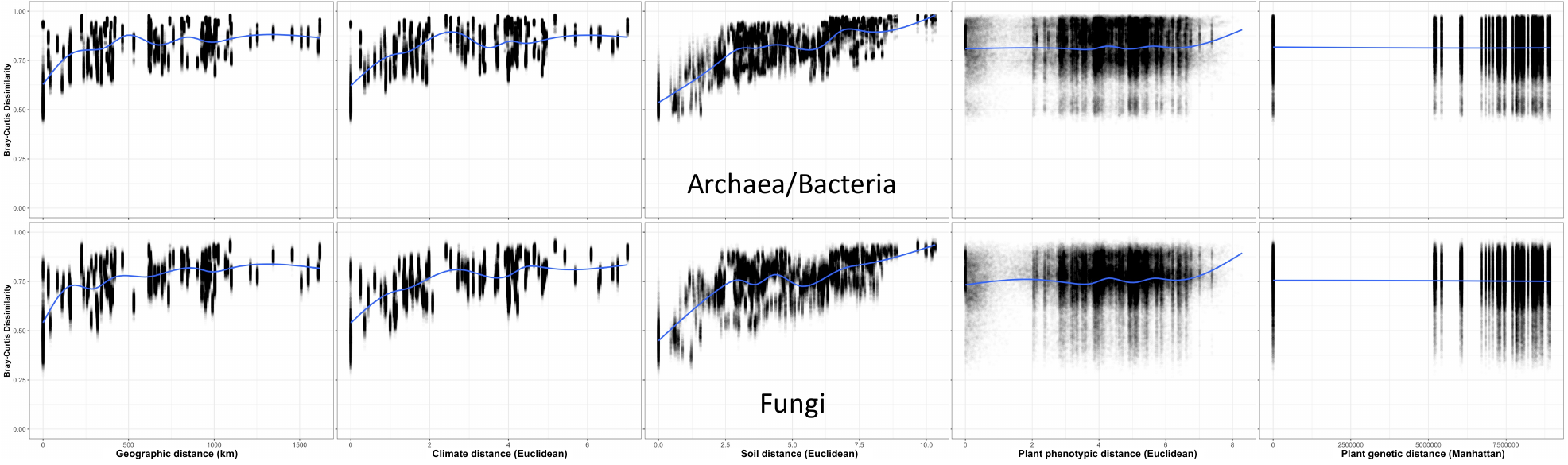

Figure S3. Correlations between (from left to right) geographic, climate, soil, plant phenotypic, and plant genetic distance and archaeal/bacterial (top row) and fungal (bottom row) Bray-Curtis dissimilarity. For statistical results, see Table 3.

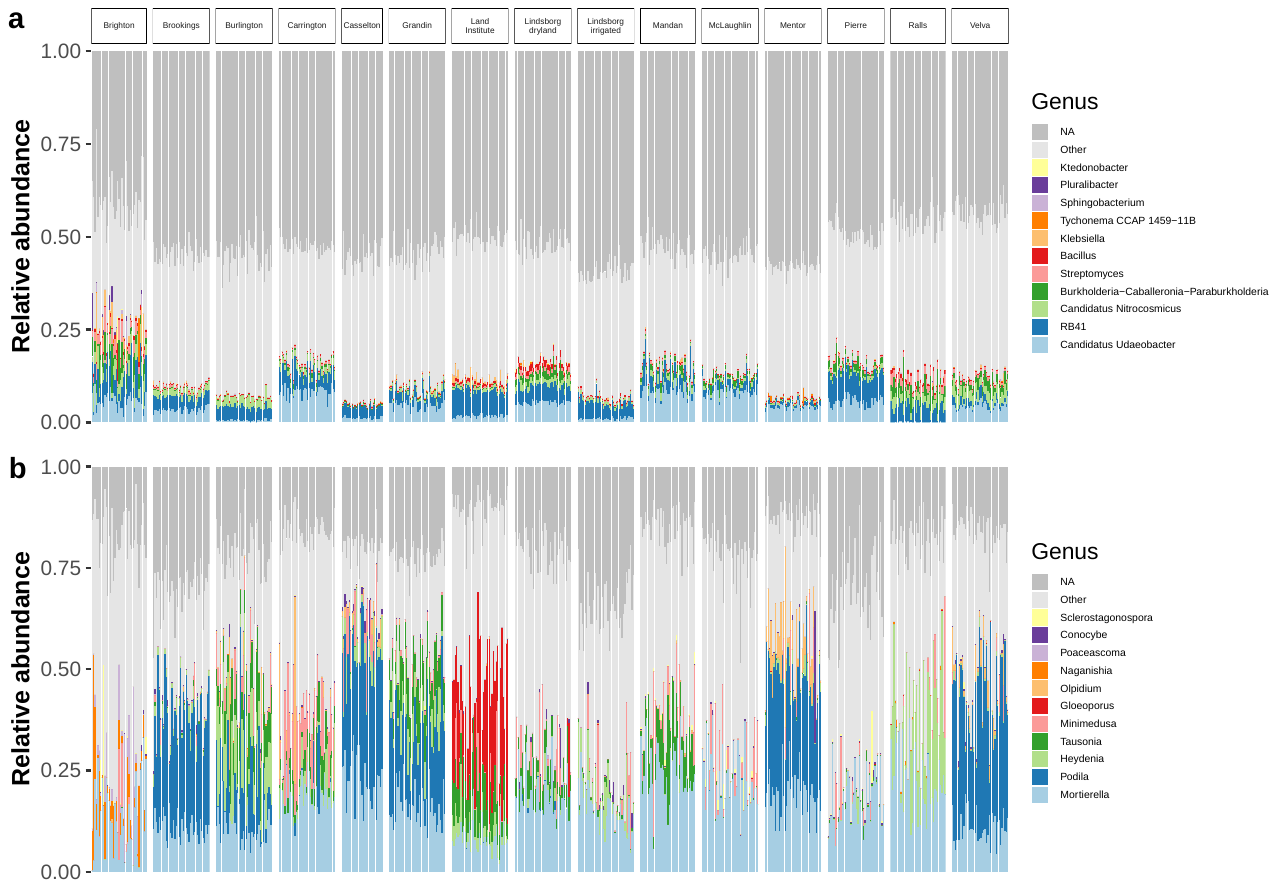

Figure S4. Top 12 a) prokaryotic genera and b) fungal genera in all rarefied samples (n = 586). All other genera not in the top 12 are aggregated into the “Other category”, while taxa unassigned at the genus level are labeled “NA”. Genera are sorted by overall abundance from bottom to top.

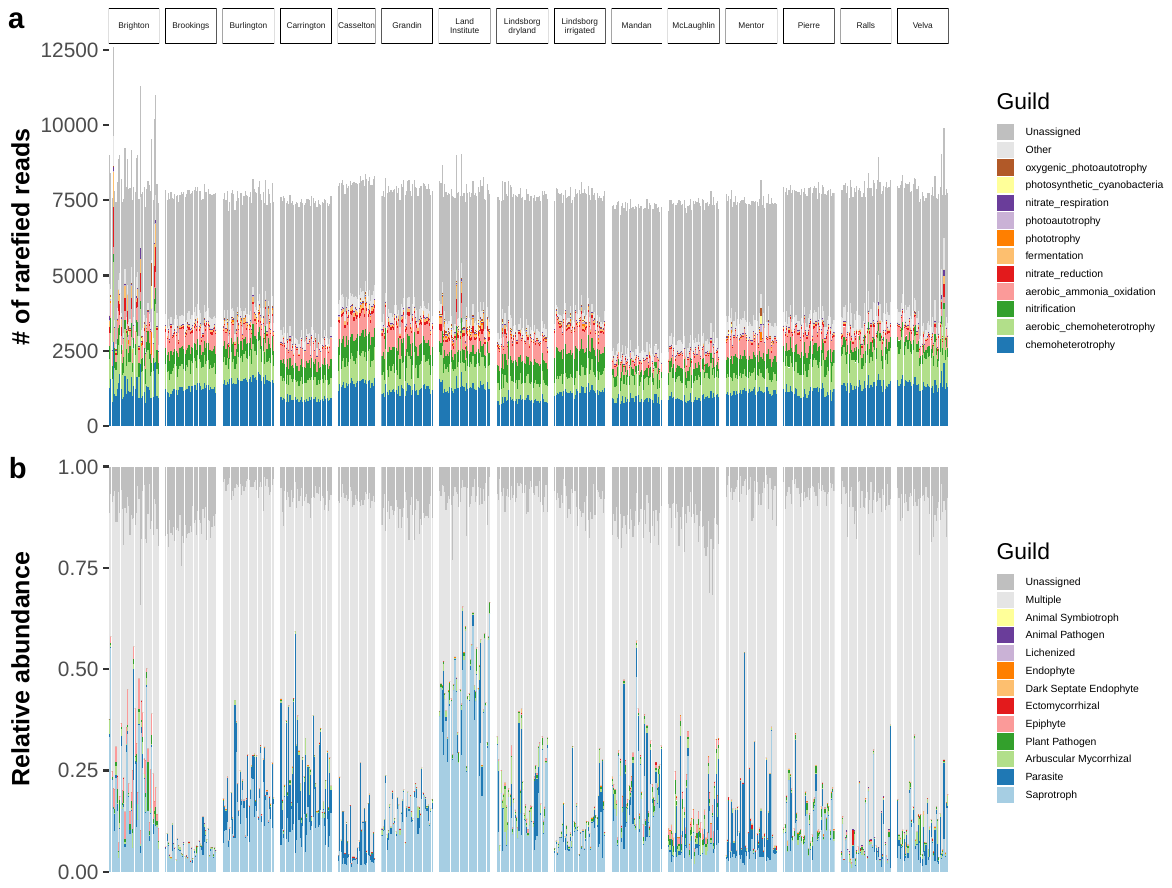

Figure S5. a) prokaryotic functional guilds according to FAPROTAX, and b) fungal functional guilds according to FUNGuild. Note that there is redundancy in the FAPROTAX assignments, as general categories such as “chemoheterotrophy” include other more detailed assignments. FUNGuild guilds included highly probable, probable, and possible assignments and were manually lumped if assigned two or more of the same category (e.g., wood saprotroph-plant saprotroph was assigned to “Saprotroph”), while guilds assigned two or more different categories were designated as “Multiple”. Dark septate endophytes were assigned as a guild based on information in the “Growth Form” output from FUNGuild.

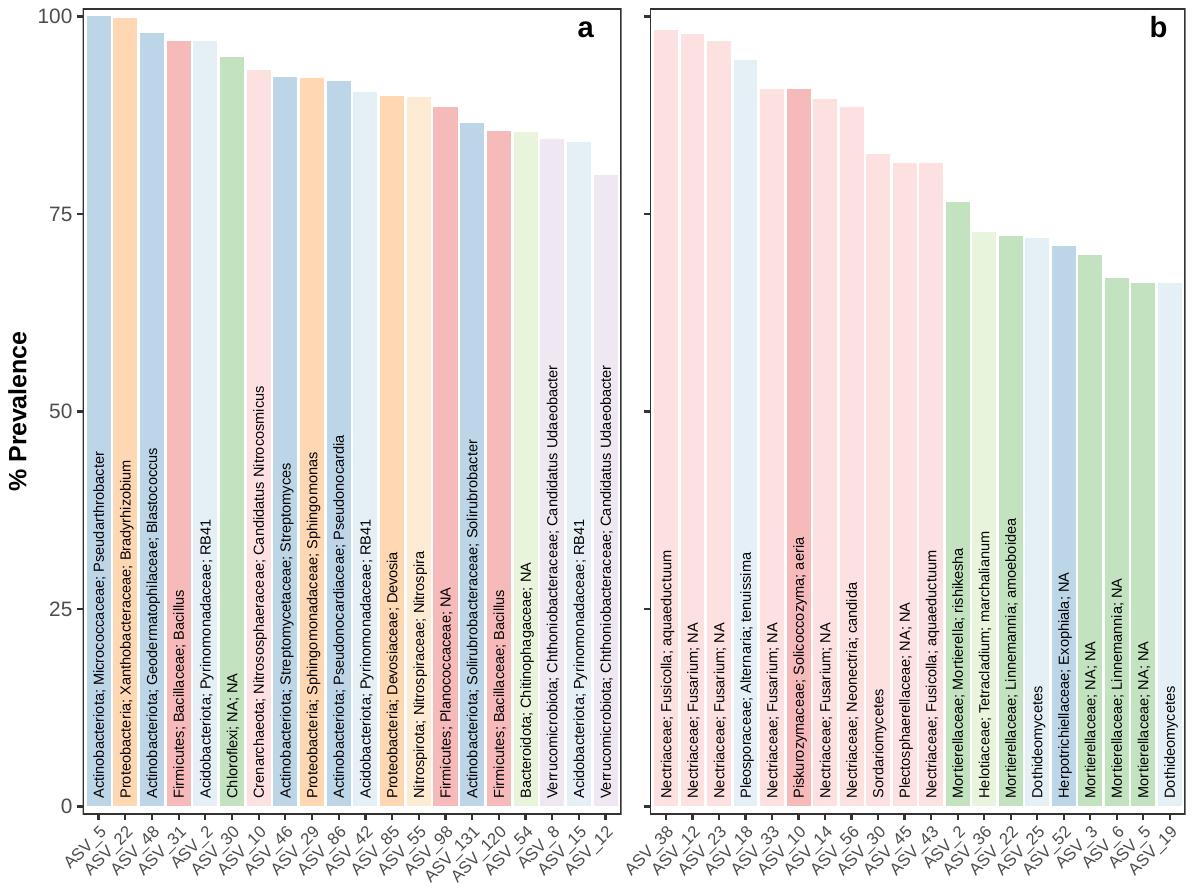

Figure S6. Top 20 most prevalent a) prokaryotic and b) fungal ASVs across the whole dataset. Bars are colored by phylum in (a) and by class in (b). Bars are labeled with Phylum; Family; Genus in (a) and with Family; Genus; Species in (b). Three ASVs with only class level taxonomic assignments in (b) are only labeled with their class.

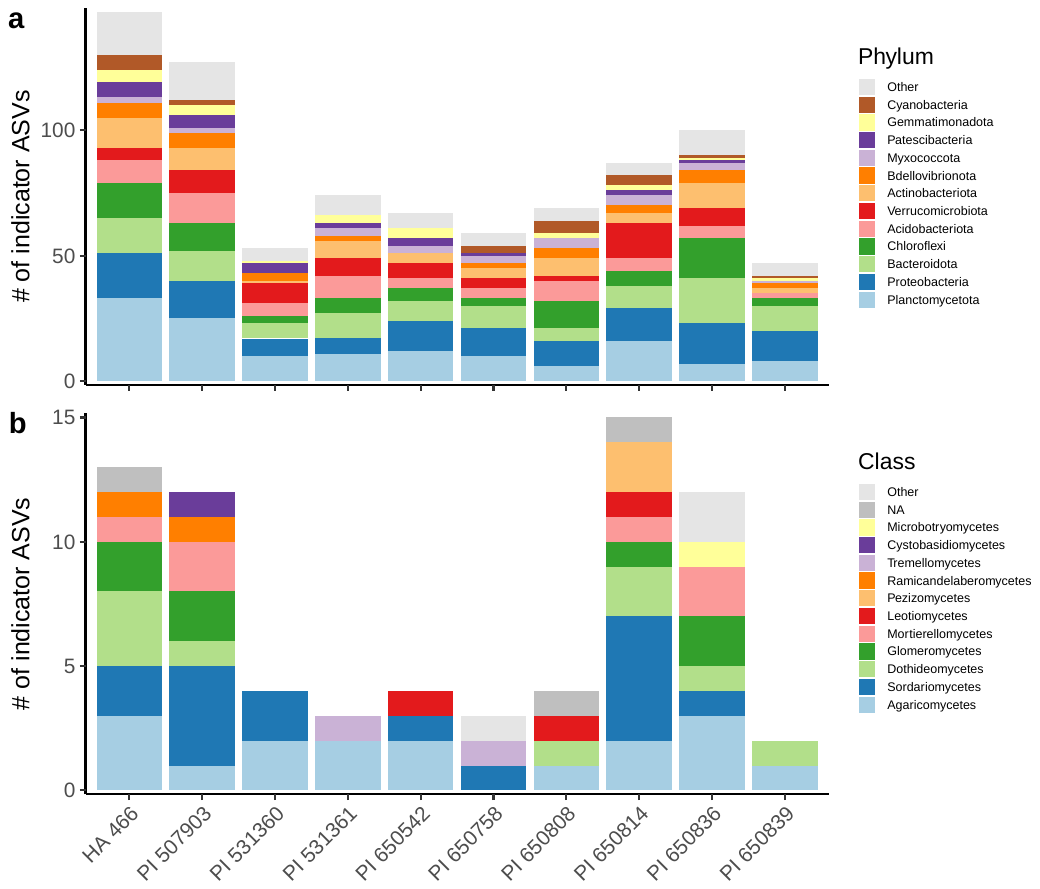
Figure S7. Number of a) prokaryotic and b) fungal genotype indicator ASVs. Bars are colored by prokaryotic phylum or fungal class. Bars and legends are sorted from bottom to top by number of ASVs in the phylum or class.

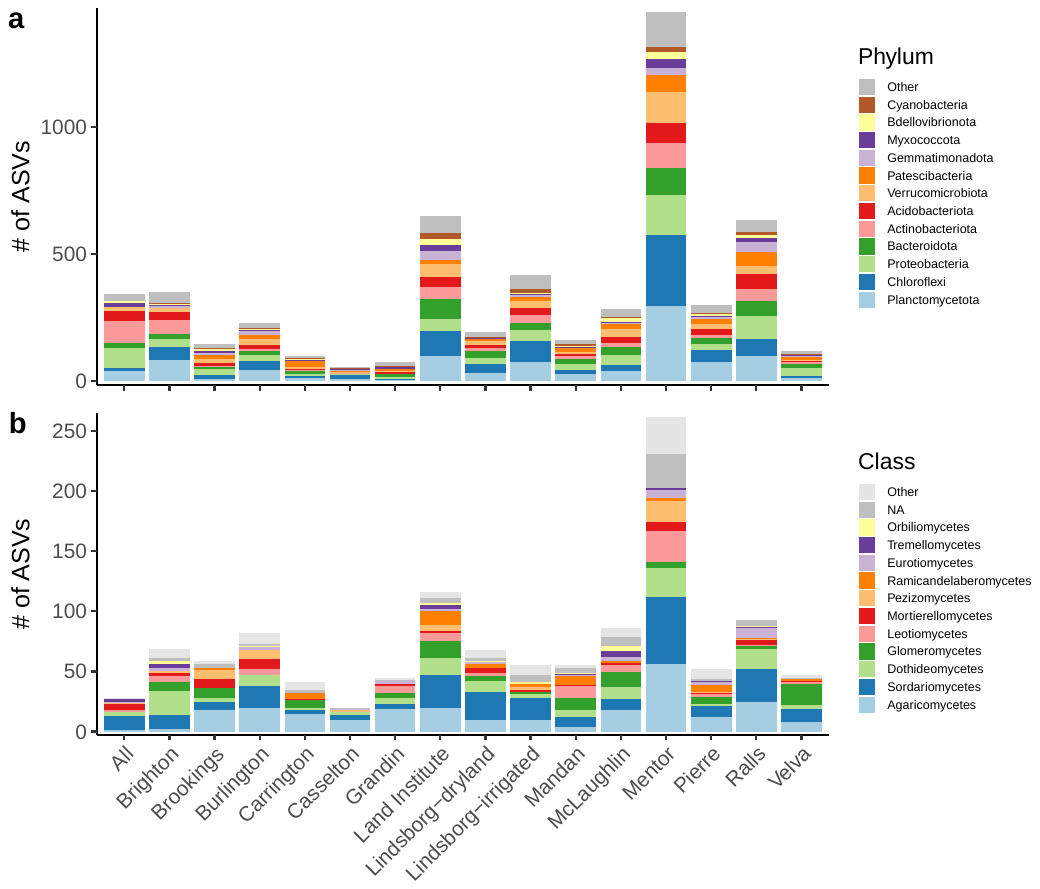
Figure S8. Number of a) prokaryotic and b) fungal ASVs present at all sites or in just one site. Bars are colored by prokaryotic phylum or fungal class. Before running the analysis, the rarefied ASV tables were further filtered such that any ASV present in only one sample in the whole dataset was removed (otherwise by definition they would be present at only one site). Therefore, ASVs present in just one site are at a minimum present in 2 samples there. Bars and legends are sorted from bottom to top by number of ASVs in the phylum or class.

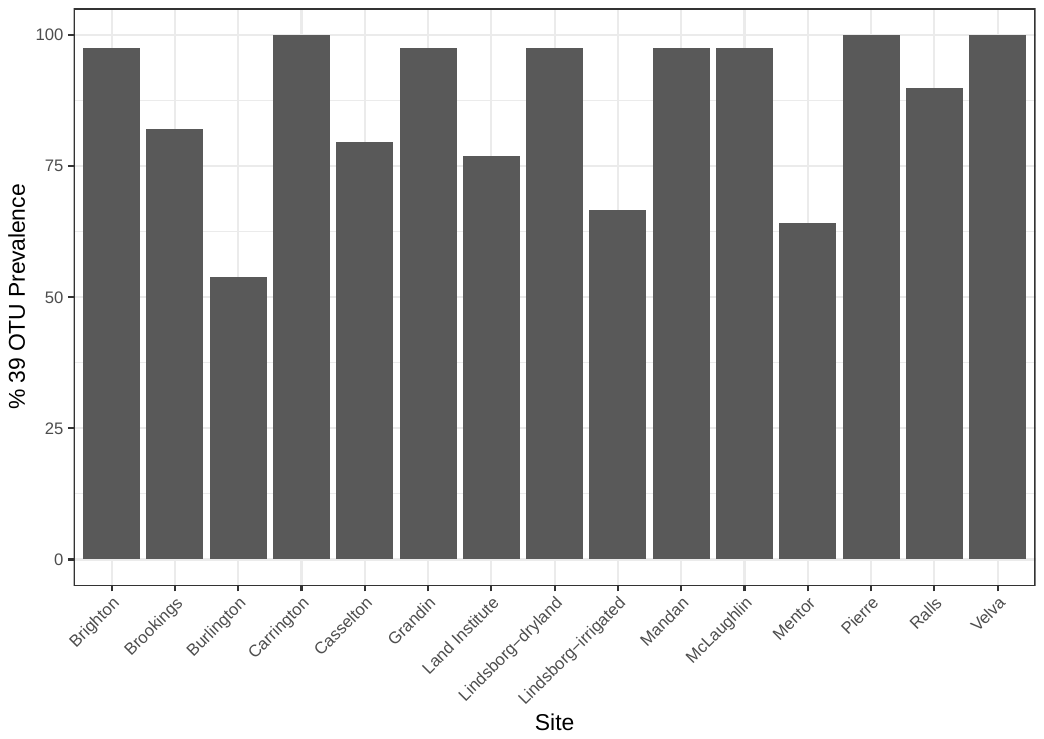

Figure S9. Percent of the 39 *Sclerotinia*-associated OTUs that were present at each site.

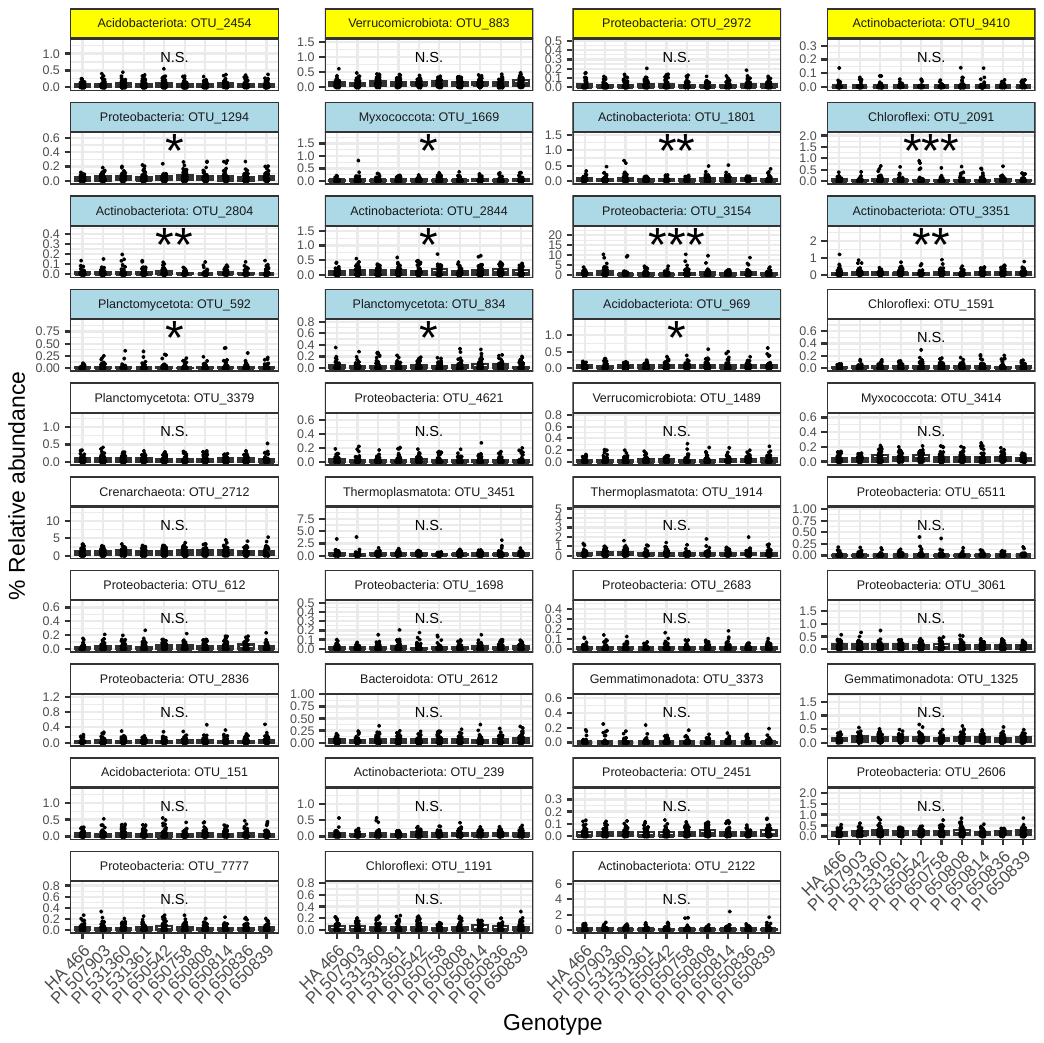
Figure S10. Effects of genotype on 39 OTUs found to be associated with *Sclerotinia* resistance in previous work at one site (Carrington). Note the y-axis scale differs among panels. Relative abundance was significantly affected by genotype for 11 of 39 OTUs (blue strips, *** = p < 0.001, ** p < 0.01, * p < 0.05). N.S. = not significant. The top row of OTUs highlighted in yellow correspond to the 4 OTUs most strongly correlated with *Sclerotinia* resistance in the previous study (Pogoda et al. 2024).
